# Spatial mitochondrial lineage tracing uncovers a premetastatic niche and microenvironment associated fate switching in osteosarcoma

**DOI:** 10.64898/2026.07.09.737611

**Authors:** Yan Xue, Zezhuo Su, Junhao Su, Alex SY Chu, Huimin Yu, Lisheng Wan, Moxian Chen, Yingxin Lin, Jason Pui Yin Cheung, Kelvin Sin Chi Cheung, Joshua WK Ho

## Abstract

Osteosarcoma progression involves complex spatiotemporal dynamics, yet the lineage relationships underlying tumor cell fate decisions remain poorly understood. Here we integrate single cell and spatial transcriptomics and mitochondrial variants-based lineage tracing to map the evolution of a late-stage, non-metastatic osteosarcoma. Single cell pseudotemporal reconstruction of malignant cells reveals a stepwise cascade of transcriptional programs that are associated with bifurcation of COL3A1⁺ progenitors into ALPL⁺ osteoblastic and THY1⁺ mesenchymal lineages. Critically, we demonstrate that the metastatic THY1⁺ mesenchymal cell fate is not governed by cell-autonomous transcriptional programs alone; this pro-tumorigenic state is associated with spatial co-localization with PLVAP⁺ dysfunctional endothelia to form a pro-metastatic niche (THY1_Endo) enriched for PTN/NOTCH signaling. To distinguish clonal ancestry from microenvironmental plasticity, we leveraged spatially resolved somatic mitochondrial variants as endogenous lineage tracers. A spatial autocorrelation filter confirmed significant clonal spatial coherence (Z = 10.07, p ≈ 4×10⁻²⁴). Phylogenetic reconstruction using a Wright-Fisher drift model coupled with a hidden Markov tree (WF-HMT) identified COL3A1⁺ cells as the ancestral tumour state with ALPL⁺ and THY1⁺ states acquired repeatedly and independently across the phylogeny. We propose a “time-space-lineage” relay model where sequential microenvironmental signals are associated with tumor cell fate and pre-metastatic niche assembly through clonal commitment. This multi-dimensional framework bridges single-cell states, spatial organization, temporal dynamics, and clonal ancestry —underscoring the power of somatic mitochondrial variants to resolve clonal history and intercept cancer evolution.

## Introduction

Osteosarcoma, the most common primary malignant bone tumor in children and adolescents, remains a formidable clinical challenge. Despite multimodal therapy, the 5-year survival rate for metastatic disease has remained below 30% for decades (Zhou et al. 2020; Zheng et al. 2024). This therapeutic ceiling reflects a fundamental gap in our understanding: how does the tumor microenvironment (TME) – a spatially complex ecosystem of malignant cells, immune infiltrates, fibroblasts, and vasculature – contribute to the transition from localized growth to metastatic competence? The TME is not merely a passive backdrop but an active participant in cancer progression, where cell–cell interactions, signaling gradients, and physical constraints collectively shape tumor evolution (Quail and Joyce 2013; Hanahan 2022). Osteosarcoma’s unique bone microenvironment adds another layer of complexity: mineralized matrix, hypoxic niches, and specialized stromal components such as cancer-associated fibroblasts contribute to chemoresistance and immune evasion through paracrine signaling (e.g., TGF-β, IL-6), extracellular matrix remodeling, and metabolic reprogramming (Cui et al. 2020; Shoaib et al. 2022). Consequently, osteosarcoma is increasingly understood not as a cell-intrinsic neoplasm but as a spatially stratified ecosystem whose immune-evasion niches choreograph progression.

Single-cell RNA sequencing (scRNA-seq) has revolutionized our appreciation of osteosarcoma intratumoral heterogeneity, revealing distinct malignant subpopulations and diverse immune landscapes. However, scRNA-seq dissociates cells from their native tissue architecture, erasing the spatial coordinates that define cellular neighborhoods, signaling distances, and physical interactions. Spatial transcriptomics (ST) bridges this gap by preserving tissue architecture while capturing transcriptome-wide expression, enabling direct mapping of molecular information onto histological structures(Ståhl et al. 2016; Maynard et al. 2021). Recent applications of ST in solid tumors have uncovered previously invisible organizational principles: spatially resolved immune evasion niches, vascular and metabolic corridors, and therapy-induced tertiary-lymphoid structures that dictate therapeutic response and resistance (Berglund et al. 2018; Moncada et al. 2020). Yet ST alone captures only a static snapshot; it cannot distinguish whether adjacent cells share a common clonal origin or have converged on similar states through convergent evolution – a distinction crucial for understanding tumor plasticity, lineage commitment, and clonal competition (Kester and van Oudenaarden 2018).

Concurrently, RNA velocity methods have emerged as powerful tools to infer transcriptional dynamics from scRNA-seq by modeling the temporal relationship between spliced and unspliced transcripts, thereby predicting future transcriptional states and uncovering the directionality of cellular transitions (La Manno et al. 2018). Cell2fate, a recent Bayesian formulation of RNA velocity, enhances this capability by decomposing transcriptional dynamics into interpretable gene expression modules – each representing a co-regulated transcriptional program – and connecting these modules to spatial tissue architecture, thereby linking temporal ordering with physical location (Aivazidis et al. 2025). In osteosarcoma, applying cell2fate could reveal whether seemingly heterogeneous tumor states represent a continuous differentiation spectrum, discrete branching events, or parallel yet convergent trajectories – each with distinct therapeutic implications.

Even with spatial and temporal dimensions integrated, however, a critical piece remains missing: lineage history. Recent technological advances have begun to address this gap by integrating CRISPR-based barcoding with spatial transcriptomics. For instance, SPACE-seq (Jia et al. 2026) and Spatio-DARLIN (Gao et al. 2026) have demonstrated the feasibility of simultaneously mapping clonal ancestry, cell state, and spatial location in mouse models of cancer and development using the DARLIN system. Importantly, while these approaches represent significant technical progress, they rely on transgenic manipulation and are therefore not directly applicable to primary human specimens. This constraint underscores the need for endogenous lineage tracers that can be retrospectively applied to clinical samples. Recent methodological advances have demonstrated that naturally occurring somatic mutations in mitochondrial DNA (mtDNA) serve as powerful endogenous lineage tracers: mtDNA is clonally inherited, accumulates mutations at a high rate, is present at high copy number per cell, and does not recombine, making it an ideal natural barcode for reconstructing cellular relationships (Ludwig et al. 2019; Lareau et al. 2021; Rodriguez-Fraticelli and Parreno 2026). Emerging methods such as MAESTER and MQuad enable enrichment and detection of informative mtDNA variants from single-cell and spatial transcriptomics libraries (Miller et al. 2022; Kwok et al. 2022; Xue et al. 2026). Moreover, by reconstructing phylogenetic trees using a Wright-Fisher drift model coupled with a hidden Markov tree (WF-HMT), one can quantitatively map clonal relatedness across space and link lineage history to transcriptional fate decisions (T. Gao et al. 2026). Importantly, because mtDNA variants are endogenous, such approaches can be directly applied to primary human tumor samples without genetic manipulation, offering a path to studying clonal evolution in patient tissues.

Here, we integrate these three analytical dimensions – space, time, and lineage – to dissect the evolution of a late-stage, non-metastatic osteosarcoma. Using ST, cell2fate RNA velocity, and mitochondrial variants-based lineage tracing on the same clinical specimen, we reconstruct a spatiotemporal evolution atlas revealing how tumor cell fate decisions and pre-metastatic niche assembly are sequentially associated with TME organization. We identify a temporally ordered cascade of 18 gene modules that execute stepwise progression: early tumor-intrinsic stress-adaptive and proliferative programs, mid-phase myeloid-driven polarization associated with COL3A1⁺ progenitor bifurcation into ALPL⁺ osteoblastic and THY1⁺ mesenchymal lineages, and late-phase formation of a spatially confined pro-metastatic niche (THY1_Endo) enriched for PTN/NOTCH signaling. Orthogonal mitochondrial lineage tracing validates COL3A1⁺ as the ancestral state, shows that the ALPL⁺ and THY1⁺ states were each acquired repeatedly and independently across the phylogeny, and identifies clade-specific mitochondrial mutations that mark terminal differentiation. Together, these findings establish a “time-space-lineage” relay model of osteosarcoma progression, bridging single-cell states, spatial organization, temporal dynamics, and clonal ancestry to provide a multi-dimensional framework for understanding and intercepting metastasis.

## Results

### Single-cell transcriptomic landscape and CNV inference delineate malignant cells from immune and non-immune microenvironments

To dissect the cellular architecture underlying osteosarcoma progression, we performed single-cell RNA sequencing (scRNA-seq) on a treatment-naïve, late-stage, non-metastatic tumor sample (Fig. 1A). After quality control, we obtained high-quality transcriptomes from 11,203 cells (Fig. S1D–F). Unbiased clustering resolved 12 distinct populations (Fig. 1B-C; Fig. S1A), reflecting the pronounced intratumoral heterogeneity characteristic of osteosarcoma.

**Fig. 1.**
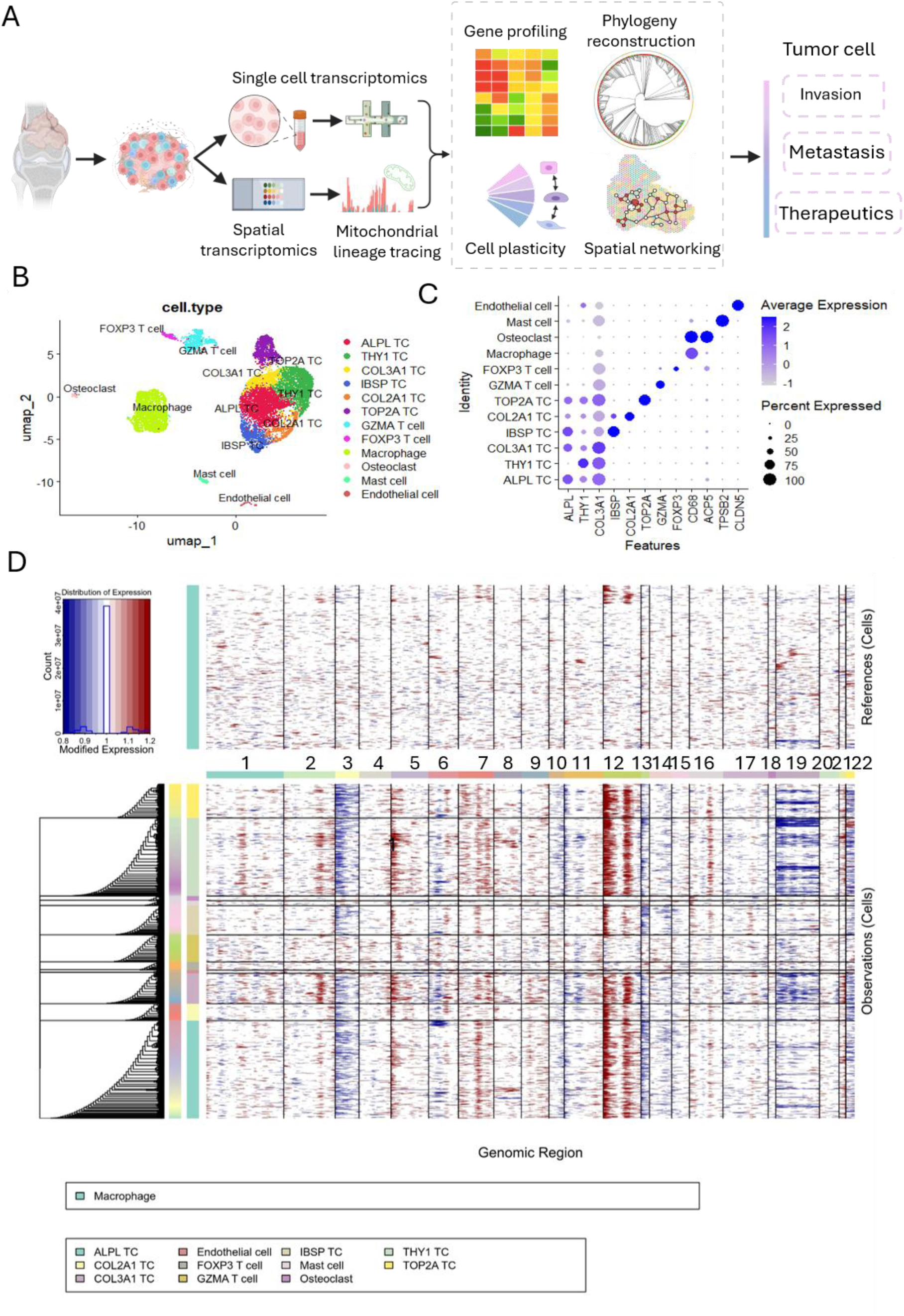
Single-cell transcriptomic analysis of the osteosarcoma lesion. (A) Schematic overview illustrating the multifaceted approach to studying osteosarcoma biology. The framework integrates single cell transcriptomics, spatial transcriptomics and mitochondrial variants enabled lineage tracing to explore the cell plasticity, cell fate, cell phylogeny and spatial networking. (B) UMAP embedding of 11,203 cells colored by unsupervised clustering, revealing 12 distinct cell populations. (C) Dot plots showing the signature gene expressions across the 12 cellular clusters. The size of dots represents the proportion of cells expressing the particular marker, and the spectrum of color indicates the mean expression levels of the markers (log1p transformed) (D) The hierarchical heatmap showing large-scale CNVs in osteosarcoma.

Among these, tumor cells constituted the dominant compartment (66.5%, 7,455 cells), followed by macrophages (22.3%, 2,498 cells) and T cells (8.2%, 915 cells) (Fig. 1B-C; Fig. S1B-C). To robustly segregate neoplastic cells from non-malignant immune and stromal populations, we integrated genome-wide copy number variation (CNV) inference with canonical cell lineage marker expression profiling to annotate malignant identity across all 12 clusters (Fig. 1C). All immune subsets (GZMA⁺ cytotoxic T cells, FOXP3⁺ Tregs, mast cells) and structural stromal cells (endothelial cells, osteoclasts) exhibited flat, near-diploid CNV profiles with negligible large-scale chromosomal gains or losses. In contrast, every transcriptionally defined tumor subpopulation carried widespread segmental amplifications on chr1, chr2, chr5, chr7, chr12 and focal deletions on chr3, chr13, hallmark genomic lesions of high-grade osteosarcoma, confirming their malignant origin (Fig. 1D).

Within the tumor compartment, we further resolved five functionally distinct subpopulations (Fig. S1B). The most prevalent subtype was characterized by high expression of *ALPL* (ALPL TCs, 22.7%), a hallmark of osteoblastic differentiation and mature bone-forming capacity (Mensali et al. 2023). The second largest cluster was defined by *THY1* expression (THY1 TCs, 18.1%), a marker frequently associated with mesenchymal stemness, cell plasticity, and therapy resistance (Wiratnaya et al. 2025). Additionally, we identified a substantial *COL3A1+* subpopulation (COL3A1 TCs, 7.1%), alongside other notable clusters such as proliferating TOP2A TCs (7.9%), IBSP TCs (6.8%), and COL2A1 TCs (3.9%) (Fig. S1B). Notably, CNV profiling revealed highly concordant large-scale chromosomal alterations between COL3A1 and THY1 subpopulations; however, THY1 is distinguished by a unique copy number profile, specifically featuring a gain at chr16 (Fig. 1D).

Beyond the tumor compartment, the non-malignant microenvironment comprised a diverse mixture of immune and stromal components. The immune landscape was highly diverse, dominated by macrophages (22.3%) and T cells (8.2%, encompassing both cytotoxic GZMA+ and regulatory FOXP3+ subsets), alongside a minor population of mast cells (1.2%). The non-immune stromal and structural compartment primarily consisted of endothelial cells (0.7%) and osteoclasts (1.1%) (Fig. S1C).

The coexistence of these transcriptionally distinct subtypes, particularly the osteogenic (ALPL+), stem-like/plastic (THY1+), and matrix-remodeling (COL3A1+) populations, suggests a highly dynamic tumor ecosystem. This distinct subpopulation architecture not only underscores the cellular plasticity of osteosarcoma but also lays the critical groundwork for exploring how specific subclones drive local invasion, orchestrate spatial networking with the immune microenvironment, and ultimately contribute to metastasis and therapeutic evasion.

### Pseudotemporal reconstruction of osteosarcoma uncovers a stepwise progression from an immune-primed niche to a pre-metastatic state

RNA velocity which involves inferring transcriptional dynamics from spliced and unspliced counts in scRNA-seq, has displayed notable potential to understand cellular dynamics in humans (La Manno et al. 2018). To reconstruct the tumor’s progression history, we applied cell2fate (Aivazidis et al. 2025), a Bayesian model of RNA velocity that is capable of inferring transcriptional dynamics in settings of complex changes or weak signals in rare and mature cell types, to this late-stage, non-metastatic sample. Pseudotemporal ordering revealed a critical fate specification event among tumor cells. Visualization of tumor cells (TCs) along pseudotime showed a clear state transition: in early pseudotime (<110 hours), the population was dominated by a progenitor-like state expressing *COL3A1* and the chondroblastic differentiation marker *COL2A1*, along with a highly proliferative *TOP2A*+ subpopulation (Fig. 2A, left). As pseudotime advanced (>120 hours), this homogeneous pool differentiated into two distinct lineages—an osteoblastic-like state marked by *ALPL* and *IBSP*, and a mesenchymal state marked by *THY1* (Fig. 2A, right).

**Fig. 2.**
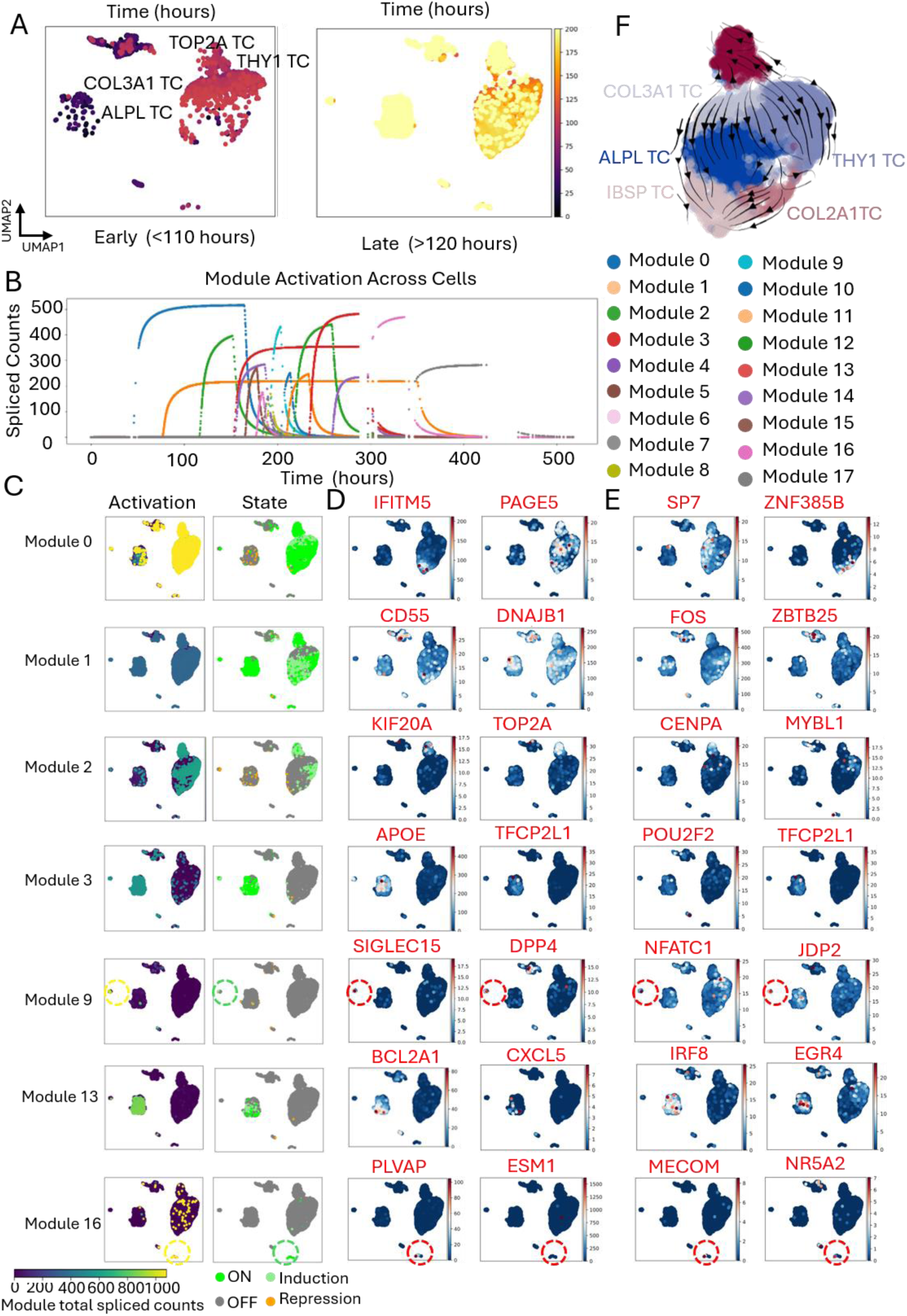
Pseudotemporal dynamics of transcriptional programs drive tumor cell fate specification and stromal remodeling in osteosarcoma. (A) Tumor cell states along pseudotime, showing early progenitor dominance and late lineage diversification. (B) Temporal activation patterns of 18 co-regulated gene modules inferred by cell2fate across pseudotime. (C) Expression patterns of key functional modules. (D) Key marker genes for each module identified based on the proportion of their transcription rate attributed to the module. (E) Key regulatory transcription factors (TFs) within each module, defined as module marker genes with known TF activity. (F) UMAP embedding of the cell2fate velocity graph, indicating transitions between tumour cell states (from COL3A1^+^ to ALPL^+^ and THY1^+^).

Cell2fate further uncovered a coordinated activation sequence of 18 gene modules that collectively elaborate the stepwise development of a pre-metastatic ecosystem (Fig. 2B-E). The trajectory was initiated by two co-emergent intrinsic programs: a canonical osteoblastic program (Module 0), defined by the master regulator SP7 and bone matrix gene IFITM5, and a concurrent stress-adaptive program (Module 1), characterized by the immune checkpoint CD55 and chaperone DNAJB1. Immediately following this foundation, a potent proliferative wave (Module 2) ensued, driven by the pro-mitotic transcription factor MYBL1 and executors such as KIF20A and TOP2A, representing a primary subpopulation targetable by DNA-damaging chemotherapy.

Subsequent to this tumor-intrinsic expansion, a coordinated immune and stromal response unfolded in mid-pseudotime. This included the sequential recruitment and functional polarization of myeloid cells: initial APOE⁺ immunosuppressive macrophages (Module 3, regulated by POU2F2) were followed by pro-inflammatory, apoptosis-resistant macrophages (Module 13, marked by CXCL5⁺ and BCL2A1⁺) that drive neutrophil recruitment and osteolysis. Concurrently, SIGLEC15⁺ osteoclasts (Module 9, driven by NFATC1) were activated, establishing a dual immune-gatekeeping role through checkpoint inhibition and chemokine degradation. This triad of macrophage/osteoclast modules (M3, M13, M9) delineates a non-linear but coherent trajectory of myeloid education from recruitment to pro-tumor inflammation and tissue remodeling (Fig. 2B–E).

The trajectory culminated in late pseudotime with the establishment of a terminal, multifaceted pre-metastatic niche. This phase was characterized by the convergence of coordinated programs, beginning with metabolic immunosuppression driven by iron-sequestering macrophages (Module 12, defined by HAMP⁺ and EGR2⁺), which create a nutrient-restricted, lymphocyte-suppressive environment. Concurrently, vascular dysfunction emerged, marked by PLVAP⁺ endothelium (Module 4/16, regulated by MECOM), facilitating aberrant permeability and likely tumor cell intravasation. These were accompanied by the activation of pro-invasive signaling programs. Notably, the macrophage phenotype displayed a functional progression, transitioning from an iron-retentive, metabolically suppressive state (Module 12) in mid-pseudotime toward a pro-inflammatory state (Module 13, CXCL5⁺) in late-pseudotime (Fig. 2B–E). This shift suggests a dynamic therapeutic target: early interventions could aim to reverse metabolic immune suppression (targeting M12), whereas late-stage strategies may need to focus on mitigating inflammation-driven tissue damage and vascular dysfunction (targeting M13 and associated vasculature programs).

Cell2fate velocity graph UMAP embedding demonstrated the high plasticity of COL3A1 TC (Fig. 2F). The velocity streamlines revealed a dynamic bifurcation originating from the early COL3A1⁺ progenitor state. Strong, concerted flows diverged into two distinct trajectories: one directed toward the ALPL⁺/IBSP⁺ mature osteoblastic branch, and the other toward the THY1⁺ mesenchymal branch. Notably, the most pronounced velocity vectors were localized at the putative branch point in mid-pseudotime, temporally coinciding with the peak activity of the macrophage-driven remodeling modules (M3, M9, M13) (Fig. S2). This alignment suggests that the tumor cell fate decision is not stochastic but is actively shaped by the evolving pressures of the myeloid-dominated microenvironment.

### Spatial transcriptomics validates the bifurcating tumor lineage and uncovers an EMT-driven metabolic switch

Tumor architecture is key to explaining the variety of observed genetic patterns (Noble et al. 2021). To delineate how spatial architecture governs tumor evolution and metastatic potential, we performed spatial transcriptomics on the same osteosarcoma sample using the 10x Visium platform (Fig. 1A and S1G–H). Using stlearn (Pham et al. 2023), which integrates gene expression similarity with spatial proximity, we identified five major spatially coherent clusters (Cluster 0 to 4) (Fig. 3A). Cell type deconvolution (Cell2location) revealed heterogeneous compositions, including multiple tumor states (ALPL, COL3A1, THY1, IBSP, TOP2A) and microenvironmental components (macrophages, FOXP3⁺ T cells, GZMA⁺ T cells, mast cells, endothelial cells) (Fig. 3B). Although both Cluster 0 and Cluster 1 were enriched in COL3A1⁺ cells, they differed in their THY1 and ALPL expression: THY1 was expressed in both clusters but was highest in Cluster 4, whereas ALPL expression was higher in Cluster 1 than in Cluster 0 (Fig. 3C). UMAP projection positioned COL3A1⁺ cells between THY1⁺ and ALPL⁺ branches, corroborating the bipotent progenitor state identified by RNA velocity (Fig. 2F and 3D). Pseudotime-space trajectory analysis further demonstrated a clear bifurcation from COL3A1⁺ progenitors (Clusters 1 and 3) toward two fates: an ALPL⁺ osteoblastic tumor lineage (Cluster 2) and a THY1⁺ mesenchymal-like tumor lineage (Cluster 4) (Fig. 3E).

**Fig. 3.**
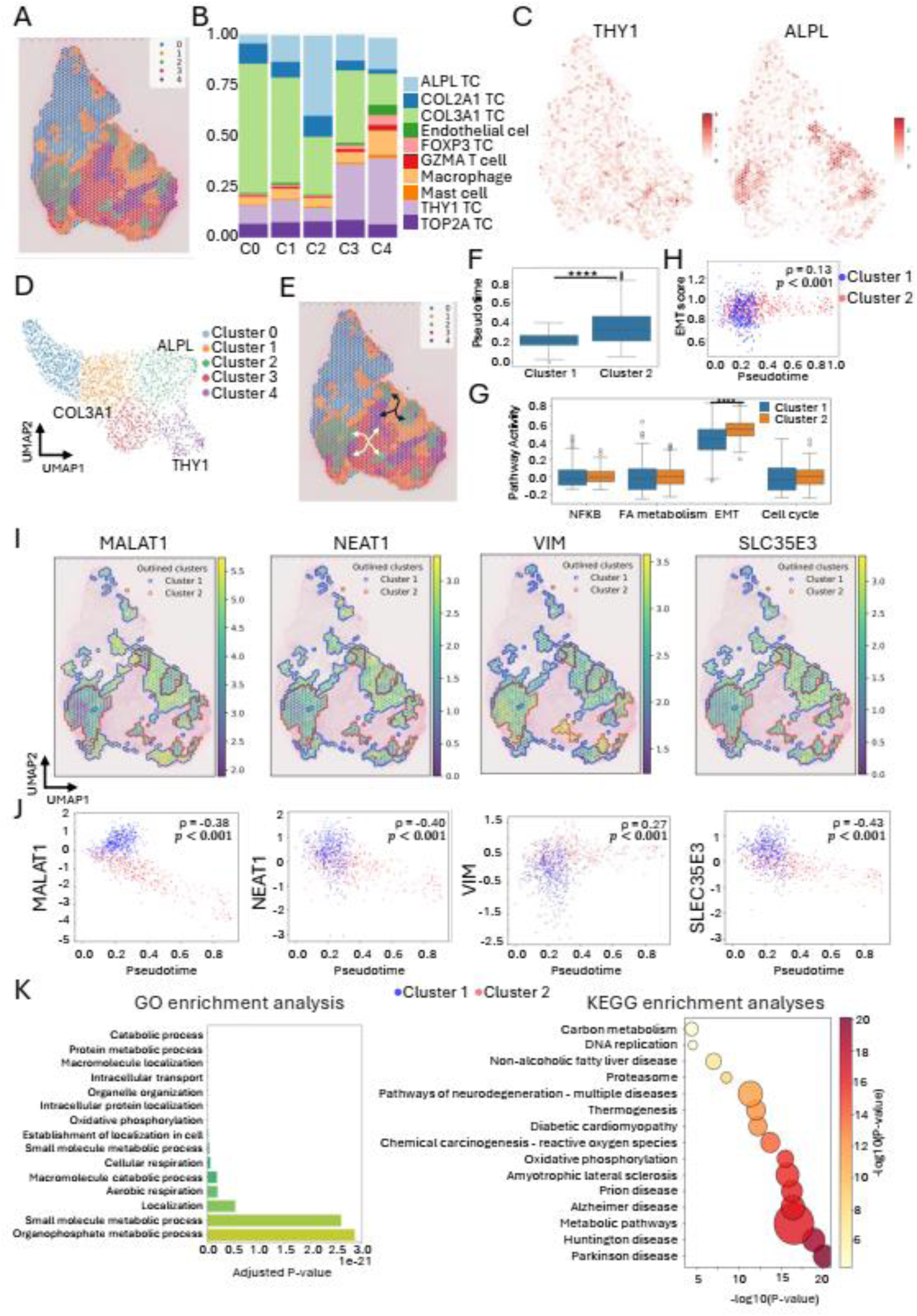
Spatial transcriptomic clustering and pseudotime trajectory reveal an EMT-driven transition from COL3A1⁺ progenitors to ALPL⁺ osteoblastic tumor cells with metabolic reprogramming. (A) Spatial clustering of Visium spots using stlearn (integrating gene expression similarity with spatial proximity) identifies five major spatially coherent clusters (0–4) in the osteosarcoma tissue section. (B) Cell type deconvolution (cell2location) reveals heterogeneous compositions, including multiple tumor states (ALPL, COL3A1, THY1, IBSP, TOP2A) and microenvironmental components (macrophages, FOXP3 ⁺ T cells, GZMA⁺ T cells, mast cells, endothelial cells). C0 to C4: Cluster 0 to Cluster 4. (C) Spatial expression maps of THY1 and ALPL. THY1 is expressed in both Clusters 0 and 1 but is highest in Cluster 4; ALPL is highest in Cluster 2 and higher in Cluster 1 than in Cluster 0. (D) UMAP projection of all cells colored by cluster identity positions COL3A1⁺ cells between THY1⁺ and ALPL⁺ branches, corroborating the bipotent progenitor state identified by RNA velocity (Fig. 2F). (E) Pseudotime-space trajectory analysis demonstrates a clear bifurcation from COL3A1⁺ progenitors (Clusters 1 and 3) toward two fates: an ALPL⁺ osteoblastic lineage (Cluster 2) and a THY1⁺ mesenchymal-like lineage (Cluster 4). Arrows indicate the inferred direction of cell state progression. (F) Pseudotime distribution boxplot shows that Cluster 2 cells occupy a significantly later temporal stage than Cluster 1 (Wilcoxon test). (G) Pathway activity scoring reveals that only the EMT pathway is robustly upregulated in Cluster 2 relative to Cluster 1. (H) EMT scores increase progressively along pseudotime. Blue and red dots represent Cluster 1 and 2 respectively. (I) Spatial mapping of key transition-associated genes (MALAT1, NEAT1, VIM, SLC35E3). VIM shows strong enrichment in Cluster 2, consistent with an acquired mesenchymal/EMT phenotype, whereas MALAT1, NEAT1, and SLC35E3 are predominantly enriched in Cluster 1. Cluster boundaries are outlined. (J) Spearman correlation analysis of gene expression along pseudotime, computed on spots pooled from both Cluster 1 and Cluster 2. Blue and red dots represent spots from Cluster 1 and Cluster 2, respectively. VIM expression significantly increases (ρ = 0.27, p < 0.001), while SLC35E3 (ρ = -0.43, p < 0.001), NEAT1 (ρ = -0.40, p < 0.001), and MALAT1 (ρ = -0.38, p < 0.001) exhibit negative correlations along pseudotime. (K–L) GO biological process and KEGG pathway enrichment analyses of differentially expressed genes (Cluster 2 vs. Cluster 1) identify top terms related to metabolic processes, including oxidative phosphorylation and reactive oxygen species metabolism.

Focusing on the transition from Cluster 1 to Cluster 2, we examined temporal dynamics and pathway activities (Fig. 3F–K, Fig. S3). Cluster 2 cells occupied a significantly later pseudotime stage than Cluster 1 (Wilcoxon test: U = 39197.0, p = 1.56×10⁻^29^) (Fig. 3F). Pathway activity scoring revealed that only the EMT pathway was robustly upregulated in Cluster 2 relative to Cluster 1(p = 8.14×10⁻^23^) (Fig. 3G), and EMT scores increased progressively along pseudotime (Fig. 3H). Spatial mapping showed that VIM was enriched in the Cluster 2 region, whereas MALAT1, NEAT1 and SLC35E3 were enriched in Cluster 1 (Fig. 3I). Consistent with these spatial patterns, VIM expression increased along pseudotime (Spearman ρ = 0.27), whereas MALAT1 (ρ = −0.38), NEAT1 (ρ = −0.40) and SLC35E3 (ρ = −0.43) progressively decreased (all p < 0.001; Fig. 3J). Together, these results link EMT-related gene reprogramming to pseudotemporal progression, supporting a trajectory from a progenitor to a mesenchymal state. GO and KEGG enrichment analyses of differentially expressed genes (Cluster 2 vs. Cluster 1) identified top terms related to metabolic processes, including oxidative phosphorylation and reactive oxygen species metabolism (Fig. 3K-L). These results indicate that the transition from peripheral COL3A1⁺ progenitors to ALPL⁺ osteoblastic tumor cells is driven by EMT activation and accompanied by a metabolic rewiring towards enhanced oxidative phosphorylation, linking spatial organization with functional reprogramming in osteosarcoma progression.

### Spatial transcriptomics reveals inflammatory reprogramming associated with NF-κB signaling during the transition from COL3A1⁺ progenitors to THY1⁺ mesenchymal-like tumor cells

We next investigated the transition from Cluster 1 to Cluster 4, which gives rise to a THY1⁺ mesenchymal-like lineage (Fig. 3E, Fig. S4). Comparative analysis showed that Cluster 4 cells occupied a significantly later pseudotime stage than Cluster 1 (Wilcoxon test: U = 2081.0, p = 1.82×10⁻^86^) (Fig. 4A). In contrast to the EMT-dominated program of the Cluster 1 to 2 transition, the Cluster 1to 4 trajectory exhibited a distinct transcriptional program centered on NF-κB-mediated inflammatory and immune reprogramming (p = 1.35×10⁻^5^) (Fig. 4B). Along pseudotime, both NFKB1 expression and the overall inflammatory score demonstrated strong positive correlations with progression (Spearman ρ = 0.24, p = 5.58×10⁻^10^ for NFKB1; ρ = 0.37, p = 2.31×10^⁻²3^ for inflammatory score), indicating a gradual and sustained activation of NF-κB signaling and inflammatory responses (Fig. 4C-D).

**Fig. 4.**
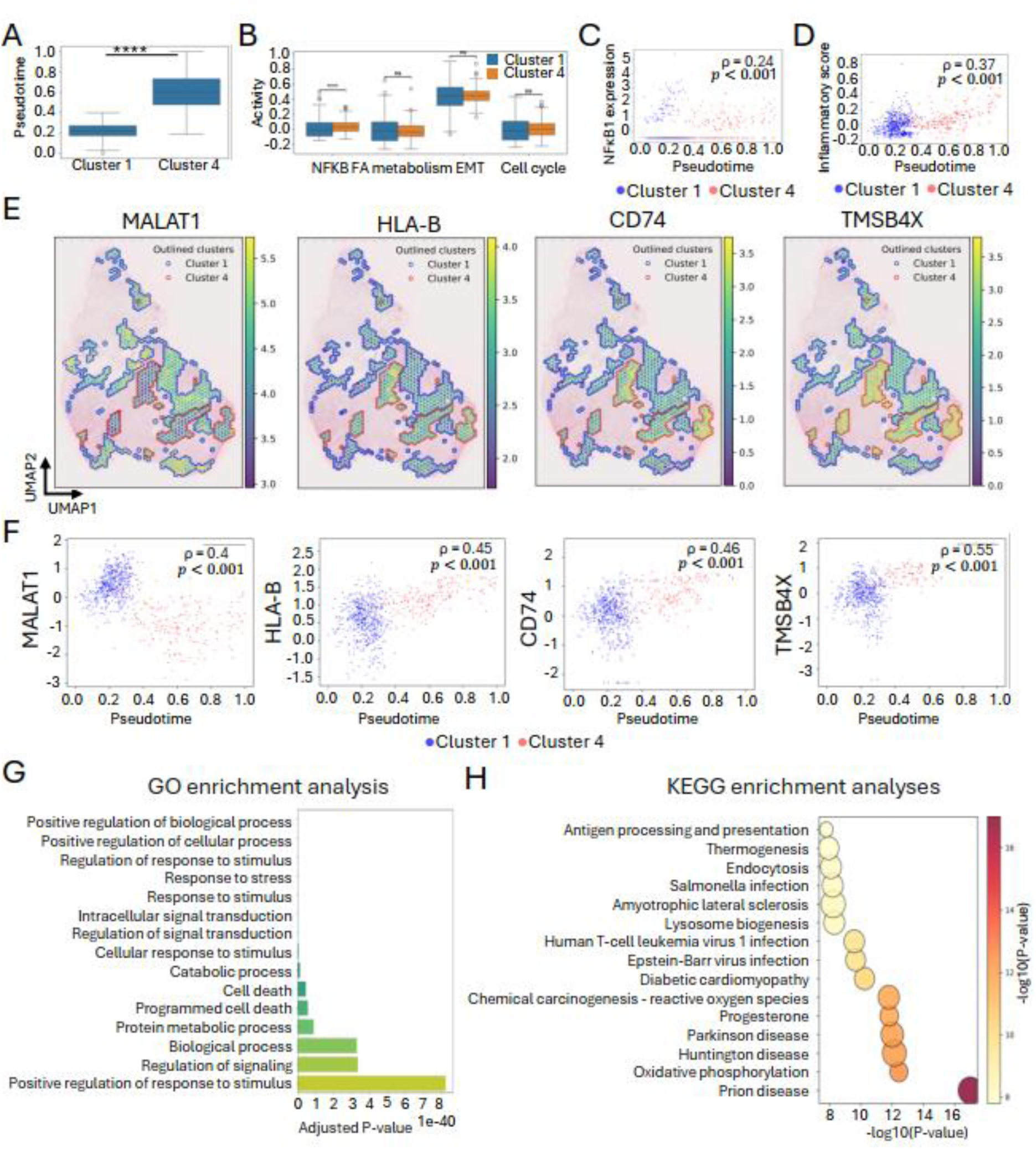
NF-κB-driven inflammatory reprogramming during the transition from COL3A1⁺ progenitors (Cluster 1) to THY1⁺ mesenchymal-like tumor cells (Cluster 4). (A) Pseudotime analysis showing that Cluster 4 cells occupy significantly later pseudotime stages compared to Cluster 1(Wilcoxon test: U = 2081.0, p = 1.82×10⁻86) . (B) Transcriptional programs along the Cluster 1 to 4 trajectory highlight NFκB-mediated inflammatory and immune reprogramming (p = 1.35×10⁻5) , distinct from the EMT program seen in the Cluster 1to 2 transition. (C, D) Spearman correlation analysis of spots pooled from both clusters along pseudotime. Along pseudotime, both NFκB1 expression (C) and an overall inflammatory score (D) exhibit strong positive correlations with progression (Spearman ρ = 0.24 and ρ = 0.37, respectively; p = 5.58×10⁻10 for NFκB1 and p = 2.31×10⁻²³ for inflammatory score). Blue and red dots represent Cluster 1 and Cluster 4 respectively. (E) Spatial mapping reveals enrichment of NF-κB-related genes (HLA-B, CD74, TMSB4X) in Cluster 4, whereas MALAT1 is enriched in Cluster 1. Cluster boundaries are outlined. (F) Spearman correlation analysis of spots pooled from both clusters along pseudotime. Pseudotemporal expression profiling confirms significant downregulation of MALAT1 along the trajectory (ρ = –0.40, p < 0.001) and positive correlations for HLA-B (ρ = 0.45), CD74 (ρ = 0.46), and TMSB4X (ρ = 0.55) (all p < 0.001). Blue and red dots represent Cluster 1 and 4 respectively. (G, H) GO (G) and KEGG (H) enrichment analyses of differentially expressed genes between Cluster 4 and Cluster 1 identify categories related to stress response, programmed cell death, oxidative phosphorylation, antigen processing/presentation, phagocytosis, endocytosis, and infection-related pathways (e.g., Human T-cell leukemia virus 1 infection), consistent with NF-κB-driven inflammatory and immune modulatory reprogramming.

Spatial mapping revealed that NF-κB-related genes (HLA-B, CD74, TMSB4X) were enriched in Cluster 4, whereas MALAT1 was enriched in Cluster 1 (Fig. 4E). Pseudotemporal expression profiling confirmed that MALAT1 was significantly downregulated along the trajectory (Spearman ρ = -0.40, p < 0.001), while HLA-B, CD74, and TMSB4X exhibited positive correlations with pseudotime (ρ = 0.45, 0.46, and 0.55, respectively; all p < 0.001) (Fig. 4F). Moreover, the inflammatory score was significantly higher in the THY1⁺ cluster (Cluster 4) than in the COL3A1⁺ cluster (Cluster 1) (Fig. S4E).

To further elucidate the biological functions underlying this transition, we performed GO and KEGG enrichment analyses on differentially expressed genes between Cluster 4 and Cluster 1 (Fig. 4G-H). GO analysis identified broad regulatory and stress-response categories, including “positive regulation of biological process,” “response to stress,” and “programmed cell death,” indicating a global shift in cellular homeostasis and stress adaptation. The KEGG enrichment analysis corroborated our previous findings, showing that Cluster 4 is characterized by strong enrichment of “Oxidative phosphorylation” and “Chemical carcinogenesis – reactive oxygen species” (consistent with metabolic rewiring), as well as “Antigen processing and presentation”, “Phagosome”, and “Endocytosis” pathways. Importantly, the presence of infection-related pathways such as “Human T-cell leukemia virus 1 infection” (a known NF-κB activator) further supports the NF-κB-driven inflammatory and immune-modulatory phenotype of Cluster 4 observed in our spatial and pseudotime analyses. Collectively, these data indicate that as COL3A1⁺ progenitors transition into the THY1⁺ mesenchymal-like state, they undergo NF-κB-driven inflammatory reprogramming coupled with metabolic changes and downregulation of antigen-presenting machinery, a mechanism that may facilitate immune evasion within the osteosarcoma microenvironment.

Analysis of an additional COL3A1⁺ progenitor cluster (Cluster 3) confirmed convergent THY1⁺ fate specification, albeit through a distinct transcriptional mechanism (Fig. S5). Having established that the THY1⁺ lineage is defined by a distinct NF-κB-driven inflammatory program—in contrast to the metabolic/EMT program of the ALPL⁺ branch—we next investigated whether this THY1⁺ state engages in specialized spatial interactions with other microenvironmental components.

### Spatial transcriptomics localizes a pro-metastatic signaling hub between THY1⁺ tumor cells and endothelial cells

To specifically resolve malignant cell states within their spatial context, we applied SpaCET, a constrained regression model designed for tumor microenvironment deconvolution (Ru et al. 2023). Interestingly, SpaCET identified a high density of malignant cells at the tissue bottom (Fig. 5A, left), which visually overlapped with regions enriched for *COL3A1*⁺, *ALPL*⁺, and *THY1*⁺ TCs as determined by cell2location (Fig. 3B,C). More importantly, SpaCET further stratified malignant cells into two hierarchical states: a less aggressive “Malignant State A” and a more aggressive “Malignant State B” (Fig. 5A, middle and right). The spatial distribution of Malignant State B showed striking overlap with the *THY1*⁺ TC patches and adjacent endothelial regions, providing independent computational validation of the aggressive THY1_Endo niche (Fig. 5A). This convergence of two distinct analytical pipelines strongly supports the biological relevance of this niche.

**Fig. 5.**
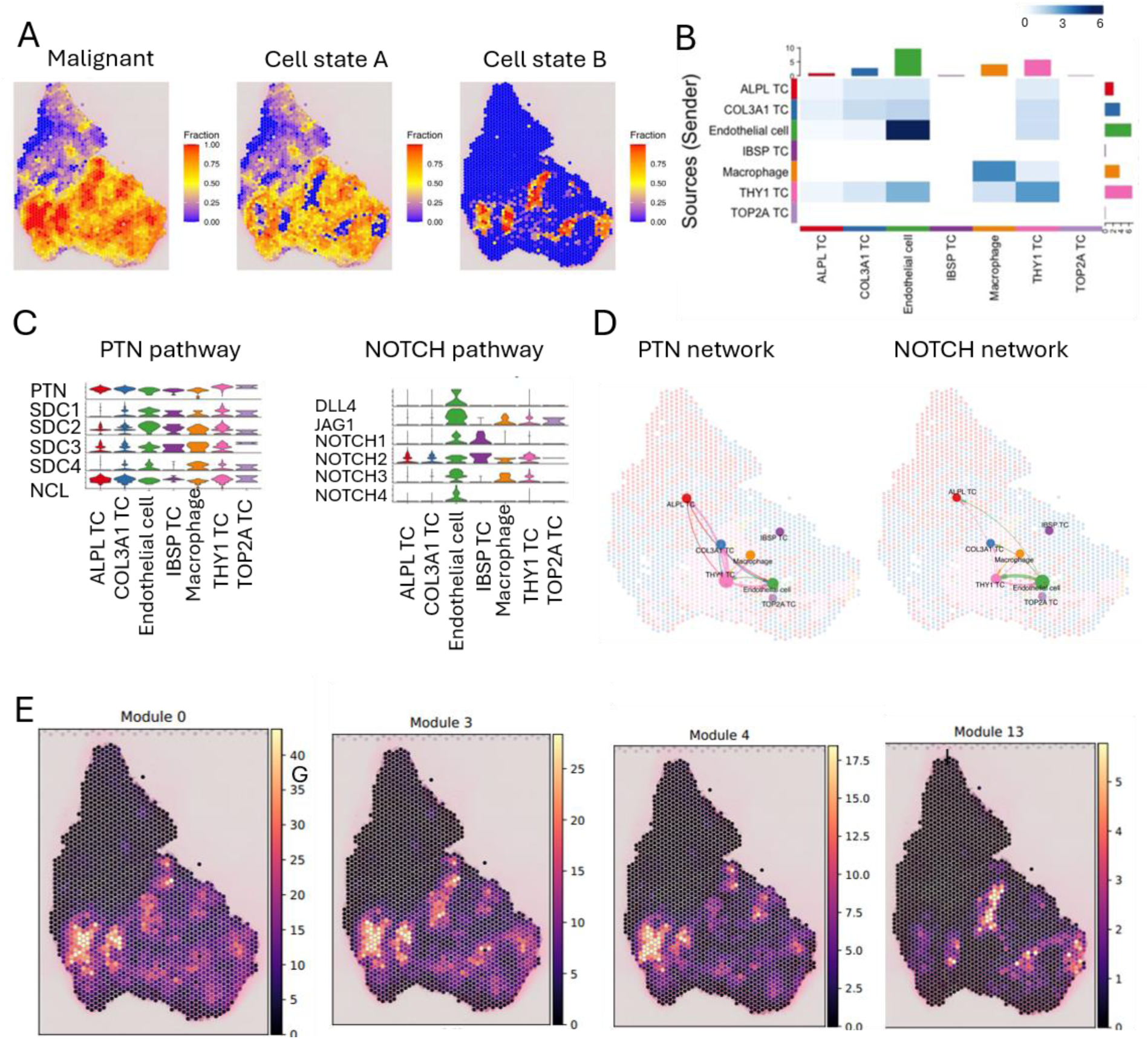
Spatial mapping of osteosarcoma microenvironment reveals PTN and NOTCH signaling mediated pro-metastasis within THY1_Endo niches. **(A)** Spatial transcriptomics analysis revealed malignant and its sub type cell states. **(B)** Spatial cell-cell interaction showing strong interaction signals between THY1 and Endo cells. **(C)** The expression of genes involved in the PTN and NOTCH pathways across the cellular composition of the THY1_Endo niches. **(D)** Spatial mapping confirms the localized interplay of the PTN and NOTCH pathways, with line thickness and vertex size representing interaction strengths. **(E)** Spatial projection of representative RNA velocity modules identified by cell2fate onto the tissue architecture, connecting spatial organization with transcriptional temporal dynamics.

To understand how physical interactions and molecular communication among different cell types regulate and coordinate tumor progression, we performed cell–cell interaction (CCI) analysis using CellChat with the ’trimean’ parameter, which effectively filters out weak (noise) interactions (Jin et al. 2025). The resulting communication network was highly specific (Fig. 5B). IBSP⁺ and TOP2A⁺ TCs showed no significant interactions with any other cell type, likely because both subtypes had low spatial transcriptomics spot counts and were shielded by the remaining five cell types. Macrophages exhibited selective, strong signaling exclusively with THY1⁺ TCs rather than with other tumor subtypes, suggesting a tailored myeloid–tumor crosstalk within this niche. Most notably, the strongest overall interaction—in both strength and number of pathways—was between THY1⁺ TCs and endothelial cells, confirming the THY1_Endo niche as a primary signaling hub within the tumor ecosystem (Fig. 5B).

Pathway analysis of this niche revealed significant enrichment for PTN (pleiotrophin) and NOTCH signaling (Fig. 5C), both critically implicated in angiogenesis, stemness maintenance, and metastatic progression. Spatial mapping confirmed the localized interplay of these pathways within the niche architecture (Fig. 5D), indicating a spatially confined, self-reinforcing signaling circuit that may drive a pro-metastatic phenotype. In summary, spatial transcriptomics transforms our view of the osteosarcoma microenvironment from a static cellular map into a dynamic decision-making landscape. Here, spatial coordinates are associated with clonal fate: THY1⁺ cells may derive their pro-metastatic identity not from cell-intrinsic programs alone, but from a spatially contingent interaction with PLVAP⁺ endothelium that confers signaling privileges unavailable to distal cells. This positions the THY1_Endo unit as a spatial dependency—a physical vulnerability that, if disrupted, could dismantle the metastatic cascade at its architectural root. The THY1_Endo niche represents a tangible spatial target whose molecular definition enables the development of precise diagnostic and therapeutic strategies aimed at intercepting the metastatic cascade.

Consistent with this spatial niche architecture, projection of the cell2fate-defined temporal modules onto the tissue revealed that mid-phase immunosuppressive modules (APOE⁺ macrophages, SIGLEC15⁺ osteoclasts) localized precisely to the interface where COL3A1⁺ progenitors commit to the THY1⁺ fate, while late-phase modules (XCL1⁺/TIAM1⁺ migration, PLVAP⁺ dysfunctional vasculature) specifically enriched within the THY1_Endo region (Fig. 5E).

This spatiotemporal integration establishes the THY1_Endo niche as a functional hub where THY1⁺ mesenchymal-like cells interface with dysfunctional endothelium through PTN/NOTCH signaling. However, a fundamental question remains unresolved: do the THY1⁺ cells within this niche arise from a common clonal lineage that expanded and differentiated, or do they represent recurrent, independently initiated mesenchymal conversions driven by local microenvironmental cues? This distinction between clonal hardwiring and plastic induction has direct implications for whether the niche can be therapeutically targeted as a lineage-defined entity or must be disrupted through its signaling dependencies. To distinguish these possibilities, we turned to mitochondrial DNA-based lineage tracing directly on the same Visium spots. Consistent with the THY1_Endo niche identified in our specimen, independent spatial transcriptomics data from nine primary osteosarcomas (Moquin-Beaudry et al. 2026), of which eight expressed COL3A1, ALPL and THY1, showed spatial co-localization of THY1⁺ tumor cells with endothelial cells (Spearman ρ = 0.50–0.89 within each sample; PT5 shown as a representative example in Fig. S6A–D; Supplementary Table 2), and cell–cell communication analysis recovered strong THY1⁺–endothelial interactions enriched for NOTCH signaling (Fig. S6E–F). In an independent osteosarcoma single-cell cohort (GSE152048; Zhou et al. 2020), the COL3A1⁺, ALPL⁺ and THY1⁺ malignant states were likewise recovered, and trajectory inference reproduced the bifurcation of COL3A1⁺ progenitors into ALPL⁺ and THY1⁺ lineages (Fig. S6G–I).

### Spatial autocorrelation of mitochondrial variants reveals clone-level spatial coherence

To resolve whether the THY1⁺ cells within the THY1_Endo niche share a common clonal origin, we leveraged naturally occurring somatic mitochondrial mutations as endogenous lineage tracers. We enriched the mitochondrial variants from Visium cDNA with MAESTER (Miller et al. 2022). However, because each Visium spot contains 10–50 cells, we first needed to establish whether mitochondrial variant allele frequencies (VAFs) measured at this mixed resolution retain sufficient spatial clonal information for reliable lineage inference. We therefore applied a Moran’s I-based spatial autocorrelation filter to MQuad-passing variants. Of eight candidate variants in the osteosarcoma sample (1,849 spots, 7,057 hex-neighbor pairs), seven cleared Bonferroni correction (α = 0.05/8 = 0.00625; Fig. 6A). The dominant variant, 1872T>A, showed strong spatial autocorrelation (Moran’s I = 0.29, Z = 24.4; Fig. 6A). Because per-spot sequencing depth (DP) was itself spatially autocorrelated (I = 0.76, Z = 65.7; Fig. 6B), we residualized each variant’s allele frequency against total DP. All seven variants remained significant after residualization (e.g., 1872T>A: Z = 13.6; Fig. 6B), confirming that the spatial signal is not a depth-driven artefact. We note that six of the seven retained variants cluster in a 12-nt window and are all A substitutions; an indirect strand-bias analysis of this pattern is provided in Supplementary Fig. S7A. To confirm that the spatial clone signal does not depend on this potentially strand-biased window, we repeated the entire clone-calling and spatial-coherence pipeline after removing the dominant variant 1872T>A (Z = 11.0) and, more stringently, after removing the entire 1861–1872 window; in the latter case the sole surviving variant, 709G>A (a G>A transition >1,150 bp away), still produced significantly coherent clones (Z = 10.5, p < 10⁻⁴; Supplementary Fig. S7B), and 709G>A remained significant after depth residualization (Z = 8.27; Fig. 6B).

**Fig. 6.**
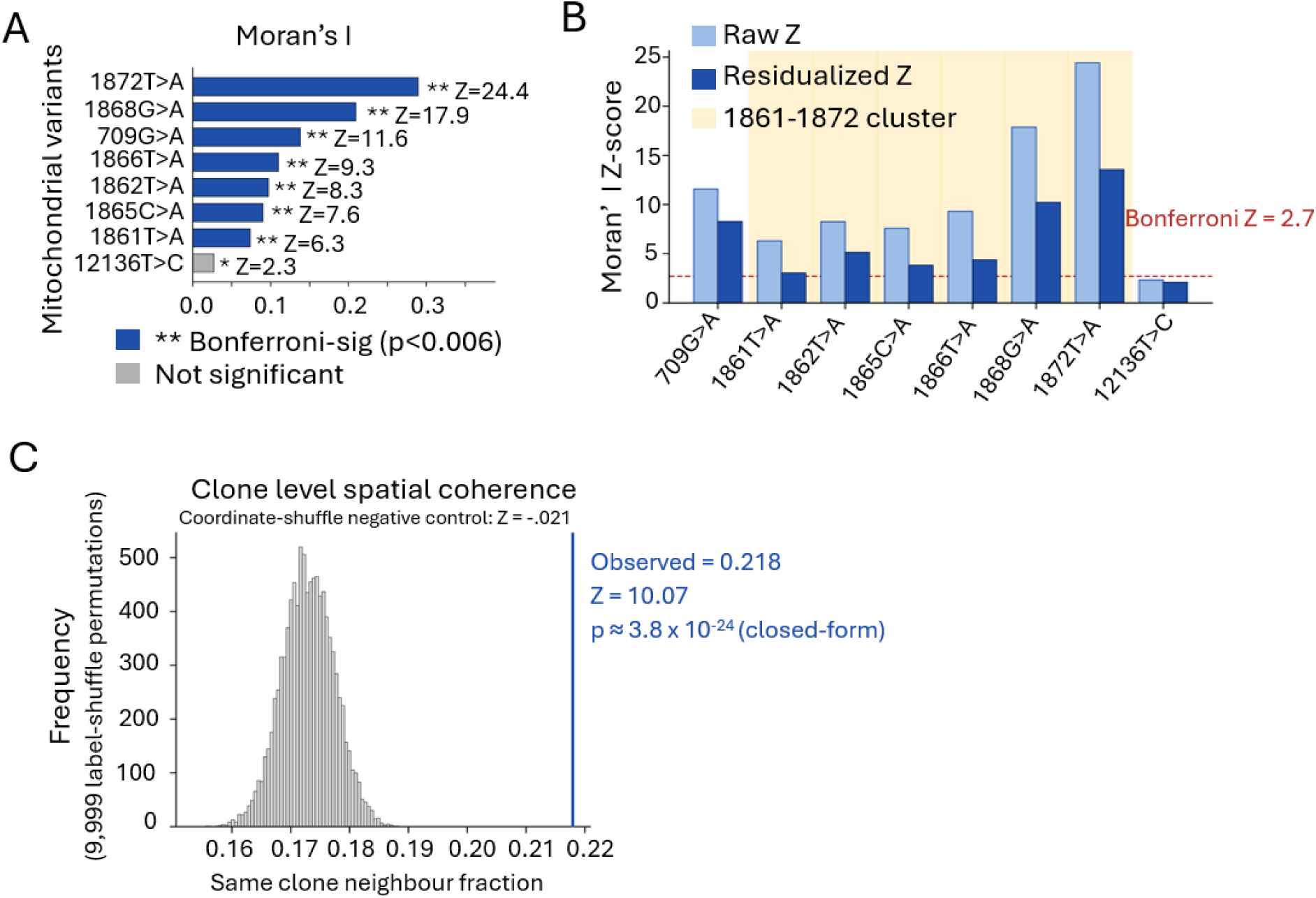
Spatial-autocorrelation-filtered mtDNA lineage analysis of Visium-MAESTER data. This analysis demonstrates that even at coarse Visium resolution (10–50 cells per spot), mtDNA variant allele frequencies (VAFs) retain sufficient spatial clonal information for rigorous lineage inference. Spatial autocorrelation filtering and clone-level coherence testing confirm that the observed clonal structure is not driven by sequencing depth or label-frequency artefacts. **(A)** Per-variant Moran’s I computed on the hex-neighbour graph (1,849 spots, 7,057 adjacent pairs), with significance assessed against 9,999 label-shuffle permutations. Seven of eight candidate variants pass Bonferroni correction (α = 0.05 / 8 = 0.00625); 12136T>C (Z = 2.3) fails and is excluded. The strongest spatial signal is 1872T>A (Moran’s I = 0.29, Z = 24.4). **(B)** Depth-residualization control. Left: comparison of raw versus depth-residualized Moran’s Z per variant. Per-spot total read depth (DP) is itself strongly autocorrelated (DP Moran’s Z = 65.7), so each variant’s allele frequency was residualized against DP by ordinary least squares. All seven Bonferroni-passing variants remained significant after residualization (e.g., 1872T>A: residualized Z = 13.6 vs. raw Z = 24.4), confirming that the spatial autocorrelation signal is not a depth-driven artefact. The substitution-class distribution of the retained variants is shown on the right. **(C)** Spatial coherence of the vireoSNP-derived six-clone partition. Histogram shows the null distribution of the same-clone hex-adjacent fraction under 9,999 label-shuffle permutations (grey). Observed fraction (0.218, blue arrow) significantly exceeds the null mean (0.173 ± 0.0045), yielding Z = 10.07 and one-sided p ≈ 4 × 10⁻²⁴ (empirical p_perm ≤ 1 × 10⁻⁴). A coordinate-shuffle negative control (inset) collapses this statistic to Z = −0.21, confirming that coherence is driven by spatial position rather than clone-label frequencies.

Using vireoSNP on the seven filtered variants yielded six spatially coherent clones. The fraction of hex-adjacent spot pairs assigned to the same clone significantly exceeded chance (observed = 0.218; null mean = 0.173 ± 0.0045; Z = 10.07, p ≈ 4×10⁻²⁴; Fig. 6C). A coordinate-shuffle negative control collapsed this statistic to Z = −0.21 (Fig. 6C, inset), confirming that coherence depends on spatial position rather than label frequencies. Thus, even at coarse Visium resolution, mitochondrial variant VAFs carry sufficient spatial clonal information to enable rigorous lineage inference.

### MitoDrift-reconstructed mitochondrial phylogeny uncovers directional COL3A1⁺ to THY1⁺ fate bias

Having validated the spatial informativeness of mitochondrial variants, we next reconstructed a high-resolution lineage tree using MitoDrift, which employs a Wright-Fisher drift model coupled with a hidden Markov tree (WF-HMT) to account for stochastic mitochondrial segregation. We collapsed low-confidence branches (τ = 0.125), yielding a confidence-refined tree inferred from 5,263 informative variants across 4,888 spots; the 1,813 spots with cell2location cell-type annotations are displayed with their predominant cell type (COL3A1⁺, ALPL⁺, THY1⁺, or other) (Fig. 7A). Posterior clade support was quantified from 16 independent MCMC chains (10,000 iterations each, 2,000-iteration burn-in): 47.9% of the 4,887 internal nodes have support ≥ 0.5, and the τ = 0.125 threshold collapses only 4 nodes (0.1%) (Figs. S8, S9). The three tumour-state clades central to our conclusions are each recovered at high frequency in a majority of the 16 chains (Fig. S10).

**Fig. 7.**
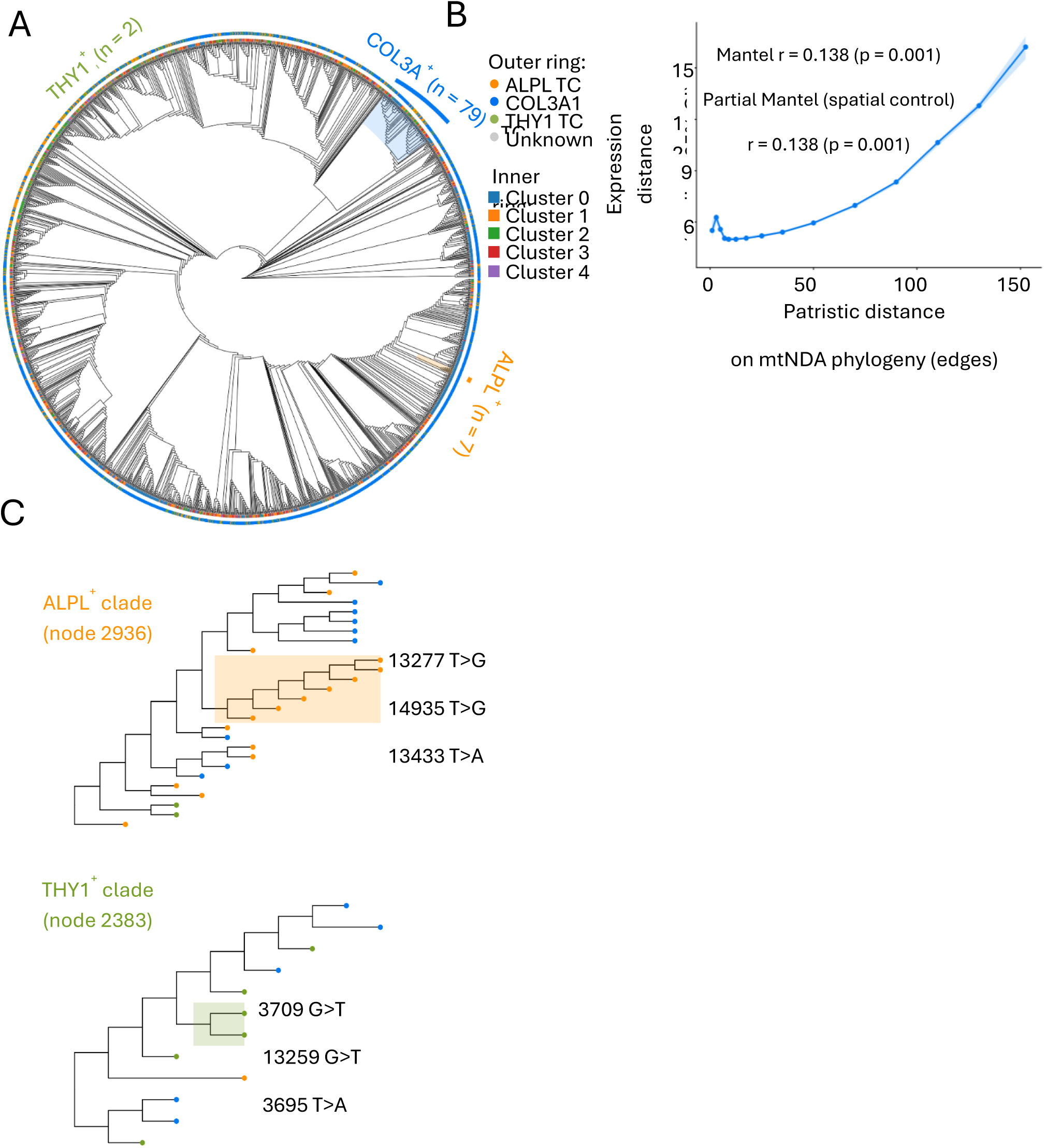
MitoDrift mitochondrial phylogeny supports an ancestral COL3A1⁺ state and repeated independent origins of ALPL⁺ and THY1⁺ states. (A) Confidence-refined MitoDrift lineage tree. Circular cladogram of the mitochondrial phylogeny inferred from 5,263 informative mitochondrial variants across 4,888 Visium-MAESTER spots (low-confidence branches collapsed at τ = 0.125), restricted to the 1,813 spots with cell2location cell-type annotations. The inner ring encodes the st-learn spatial cluster; tip points are coloured by the dominant cell type (COL3A1⁺ = blue, ALPL⁺ = orange, THY1⁺ = green, unassigned = grey). The highest-purity clade for each tumour state is highlighted and labelled: COL3A1⁺ (node 1886, n = 79, posterior support 0.557), ALPL⁺ (node 2936, n = 7, support 0.564) and THY1⁺ (node 2383, n = 2, support 0.829). Posterior clade support was computed from 16 independent MCMC chains × 10,000 iterations (2,000-iteration burn-in). (B) Lineage–state coupling is concentrated among close relatives. Median expression distance (95% CI) as a function of patristic distance on the phylogeny: expression distance is low and flat for spots sharing a recent common ancestor and rises for distantly related spots. Mantel r = 0.138 (p = 0.001); partial Mantel controlling for spatial distance r = 0.138 (p = 0.001). (C) High-purity ALPL⁺ (node 2936) and THY1⁺ (node 2383) clades shown as focused subtrees together with their immediate phylogenetic neighbourhood; the three most clade-characteristic mtDNA mutations are annotated beside each clade (ALPL⁺: 13277 T>G, 14565 T>G, 13433 T>A; THY1⁺: 3709 G>T, 13259 G>T, 3695 T>A; one-sided Fisher exact screen, Benjamini–Hochberg FDR; Supplementary Table 1). Per-spot variant allele frequencies of these mutations across the displayed spots are shown in Fig. S14.

To test lineage–state coupling, we calculated phylogenetic distance (cophenetic distance) and transcriptomic distance (Euclidean on PCA) for every spot pair. A Mantel test revealed a significant but weak positive correlation (r = 0.138, p = 0.001) that was unchanged after controlling for spatial distance (partial Mantel r = 0.138, p = 0.001; Fig. 7B), indicating that the coupling is genuine but modest at the whole-tree level. The signal is concentrated among close relatives: expression distance is low and flat for spots sharing a recent common ancestor and rises only for distantly related spots (Fig. 7B), and each spot’s closest relative on the tree is transcriptionally more similar than spatially matched null pairs (Wilcoxon p = 3.5 × 10⁻⁶; Fig. S11).

The phylogeny is rooted by construction through an ancestral dummy outgroup (see Methods), and the root state is reconstructed as COL3A1⁺ under outgroup, midpoint and burden-minimal rootings alike. Two model-free lines of evidence independently support a COL3A1⁺-first ordering: tips immediately adjacent to the root carry minimal derived-variant burden (median 2 versus 282 genome-wide; Wilcoxon p = 1.6 × 10⁻⁸), and per-spot derived-variant burden increases along the COL3A1⁺→THY1⁺→ALPL⁺ axis after controlling for sequencing depth (Fig. S12). Maximum-parsimony ancestral-state reconstruction further resolves extensive polyphyly: the THY1⁺ state is acquired independently ∼186 times (versus 41 losses) and the ALPL⁺ state ∼137 times (versus 75 losses). Consistent with repeated acquisition, Blomberg’s K (Blomberg et al. 2003) was < 0.1 for all three states (Fig. S13), and same-state sister-tip (cherry) pairs are enriched for COL3A1⁺ (301 observed versus 269 expected, p = 0.001) and ALPL⁺ (31 versus 11, p = 0.001) but not for THY1⁺ (10 versus 8.9, p = 0.39), with per-state phylogenetic sorting scores showing the same pattern (COL3A1⁺ Z = 13.2 and ALPL⁺ Z = 15.1, versus THY1⁺ Z = 1.9, p = 0.059). Thus THY1⁺ and ALPL⁺ cells arose multiple times independently from COL3A1⁺ progenitors rather than descending from single monophyletic clades.

Finally, we identified high-purity THY1⁺ and ALPL⁺ clades (nodes 2383 and 2936, purity = 1; posterior support 0.829 and 0.564), which are highlighted directly on the full tree (Fig. 7A; Fig. S15) and shown as focused subtrees together with their characteristic mtDNA mutations (Fig. 7C; Fig. S14). A systematic one-sided Fisher exact screen of all 5,263 variants against every high-purity clade (Benjamini–Hochberg FDR) found the COL3A1⁺ and ALPL⁺ clades to be supported by 372 and 100 distinct enriched variants, respectively (Supplementary Table 1), demonstrating that the inferred clades rest on many co-inherited variants rather than any single mutation. Clade-specific mitochondrial mutations showed high prevalence within these clades but were rarely detected elsewhere, indicating that these mutations accumulated prior to, rather than as a consequence of, the phenotypic switch.

Collectively, MitoDrift-inferred phylogenies demonstrate that COL3A1⁺ tumor cells act as the ancestral state, with ALPL⁺ and THY1⁺ states acquired repeatedly and independently across the phylogeny and marked by distinct mutation signatures.

## Discussion

In this study, we present an integrated spatiotemporal atlas of a late-stage, non-metastatic osteosarcoma, directly linking sequential transcriptional modules to cell fate bifurcation and pre-metastatic niche assembly. Our key finding is the identification of the THY1_Endo niche, a spatially confined hub where THY1⁺ mesenchymal tumor cells interface with PLVAP⁺ endothelial cells, as the structured endpoint of a temporally ordered cascade.

A central challenge, recently brought into sharp focus by studies in colorectal cancer, is to distinguish whether transcriptionally defined pro-metastatic states arise from transient microenvironmental plasticity or from deterministic clonal lineage locking (Buissant Des Amorie et al. 2026). In that system, the observed complete plasticity between canonical and oncofetal states left unresolved whether cell-extrinsic signals alone dictate cancer cell identity, or whether clonal ancestry imposes hidden constraints (Bulliard et al. 2026). A parallel observation comes from SPACE-seq in intrahepatic cholangiocarcinoma (iCCA), where clonally related tumor cells were found to diversify into distinct differentiation states, suggesting intratumoral phenotypic plasticity (Jia et al. 2026). However, because SPACE-seq relies on CRISPR barcodes that do not explicitly model directional transition rates, it remained unclear whether these states represent reversible fluctuations or irreversible lineage commitment—a distinction that is critical for understanding whether such heterogeneity is therapeutically targetable. To resolve this fundamental question, we leveraged somatic mitochondrial variants as natural genetic barcodes—they accumulate mutations at a high rate, are maternally inherited without recombination, and can be sequenced from the same spatial transcriptomics spots, enabling direct integration of clonal ancestry with spatial context.

Reconstructing a phylogenetic tree using a Wright-Fisher drift model coupled with a hidden Markov tree (WF-HMT), which explicitly accounts for stochastic mtDNA heteroplasmy drift during cell division, independently validated our RNA velocity inferences. The Mantel correlation between mitochondrial lineage distance and transcriptomic distance, though weak at the whole-tree level (r = 0.138, p = 0.001), is concentrated among close relatives and is not spatially confounded, indicating that recent mitochondrial ancestry predicts similar transcriptional states and supporting the biological relevance of our phylogenetic reconstruction. Ancestral-state reconstruction identified COL3A1⁺ as the root state under three rooting strategies, and derived-variant burden increased along the COL3A1⁺ to THY1⁺→ALPL⁺ axis independently of sequencing depth—model-free evidence for a COL3A1⁺-first ordering. Parsimony reconstruction further indicated that the THY1⁺ and ALPL⁺ states were each acquired repeatedly and independently (∼186 and ∼137 origins, respectively). The low Blomberg’s K values (all <0.1) further indicate that neither THY1⁺ nor ALPL⁺ cells are confined to single monophyletic clades, consistent with multiple independent transition events from ancestral COL3A1⁺ progenitors. Clade-specific mitochondrial mutations (prevalence >0.5) within high-purity THY1⁺ and ALPL⁺ clones further support that mutation accumulation preceded and may have permissively enabled the phenotypic switch.

Beyond clonal dynamics, the spatial dimension of this lineage architecture yields an unexpected insight. The significant spatial coherence of mitochondrial clones in this osteosarcoma (Z = 10.07, p ≈ 4×10⁻²⁴) is consistent with limited cell dispersal during clonal expansion, a pattern documented in normal epithelia and haematopoiesis (Lareau et al. 2021; Penter et al. 2021; Ludwig et al. 2019; Miller et al. 2022) but less characterised in solid tumors with a complex stromal architecture. Our spatial autocorrelation filter, which screens MQuad-passing variants by Moran’s I, adapts the logic of spatially variable gene detection methods such as SpatialDE (Svensson et al. 2018), SPARK (Sun et al. 2020), and SUMMIT (Bracht et al. 2025) from gene-expression surfaces to the variant-quality problem in Visium-MAESTER data.

This clonal and spatial framework gains particular significance when integrated with recent work identifying THY1⁺ cancer stem cells (CSCs) as key drivers of metastasis across multiple cancer types (Wan et al. 2026). In that paradigm, THY1⁺ CSCs exploit a pseudohypoxic state and engage in direct communication with neutrophils via THY1–Mac1 interactions to acquire pro-metastatic machinery. Our data independently corroborate and substantially extend this framework in osteosarcoma: THY1⁺ cells emerge recurrently and independently across the tumour phylogeny, yet once acquired the state is rarely reversed, and these cells co-localize with PLVAP⁺ dysfunctional endothelium into a spatially confined signaling hub (THY1_Endo) enriched for PTN/NOTCH signaling. The strong excess of THY1⁺ gains over losses in our parsimony reconstruction (186 versus 41 state changes) indicates that, once a COL3A1⁺ progenitor commits to the THY1⁺ fate, reversal is rare—distinguishing this mesenchymal-like state from a merely transient phenotype, while the spatial niche architecture reveals where this state localizes and with which signalling it is associated. This directional clonal dynamic, combined with the spatial niche architecture we describe, provides a unifying framework: THY1 marks not only a pro-metastatic, stem-like population, but one whose phenotype is both lineage-associated (a directional fate bias with rare reversal) and niche-associated (spatially restricted co-localization with PLVAP⁺ endothelium and PTN/NOTCH ligand–receptor signalling). Therapeutically, this dual dependency suggests that targeting THY1⁺ cells may require simultaneous disruption of their lineage commitment and their niche support, a combination strategy that could be more effective than targeting either axis alone.

Collectively, our spatial data argue that the THY1_Endo niche is not a passive destination for migratory tumor cells, but an active induction hub where spatial proximity may generate a signaling threshold that is associated with the metastatic program. This places spatial architecture on equal footing with genetic mutations in determining metastatic outcome, a concept that transforms the TME from a modulatory influence into a spatially encoded component of metastasis.

From a clinical perspective, the THY1_Endo niche offers a spatial biomarker for metastatic risk stratification. The temporal modularity of progression supports stage-specific intervention: early targeting of the proliferative module, mid-phase myeloid-directed agents to disrupt fate specification, and late-phase niche-disrupting combinations (PTN/NOTCH inhibition). Importantly, the mitochondrial mutations we identified in terminal clones could serve as clonal markers to track minimal residual disease or assess therapeutic response.

Several limitations should be acknowledged. Our analysis is derived from a single tumor section, and validation across independent cohorts and osteosarcoma subtypes is required. The multi-cell resolution of Visium limits direct cell-level clonal assignment; emerging single-cell spatial technologies (e.g., MERFISH, CosMx) will enable finer-resolution niche mapping. Additionally, while we controlled for depth-driven artefacts, deeper MAESTER sequencing and joint statistical models incorporating spatial smoothing would further improve clonal boundary resolution. Regarding mitochondrial variants analysis, we acknowledge that heteroplasmy thresholds, sequencing depth, and potential nuclear mitochondrial DNA segments (NUMTs) require careful filtering. Future work integrating mitochondrial lineage tracing with patient-derived organoids or xenografts could causally link specific clonal mutations to metastatic behavior.

In summary, our integrated analysis demonstrates that osteosarcoma metastasis is a spatiotemporally programmed process. By combining RNA velocity, spatial transcriptomics, and mitochondrial variant-based lineage tracing with a Wright-Fisher/HMT phylogenetic model, we provide a dual-axis (time and space) framework for understanding and therapeutically targeting the metastatic cascade. This work also exemplifies how somatic mitochondrial mutations — often considered neutral bystanders — can be harnessed as powerful endogenous recorders of clonal dynamics in cancer.

## Methods

### Patient and sample collection

The osteosarcoma sample was obtained from a 46-year-old male patient who underwent surgical resection of a primary tumor arising from the femur. The diagnosis of high-grade (Grade III) conventional osteosarcoma with chondroblastic differentiation was confirmed by two independent pathologists based on the World Health Organization classification criteria, with the presence of neoplastic osteoid and malignant cartilage. The patient had not received any chemotherapy or radiotherapy prior to surgery. Fresh tumor tissue was collected immediately after resection, transported on ice, and processed within 24 h. Written informed consent was obtained from the patient, and the study protocol was approved by the Institutional Review Board of the University of Hong Kong/Hospital Authority Hong Kong West Cluster (IRB reference number: UW 16-2036 and UW 13-576). All methods were performed in accordance with relevant guidelines and regulations.

### Tissue dissociation and single-cell suspension preparation

Fresh tumor tissue was collected immediately after surgical resection and transported on ice. The tissue was washed 2–3 times with HBSS (Gibco, Cat. No. 14025076) and minced into small nodules (approximately 1 mm³). To remove peripheral blood cell contamination, the minced tissue was washed an additional 2–3 times with HBSS. Digestion was performed using an enzyme cocktail consisting of 0.4% (w/v) collagenase II (Gibco, Cat. No. 17101015), 0.4% (w/v) hyaluronidase (Sigma, Cat. No. H3884-1G), and 0.4% (w/v) dispase II (Sigma, Cat. No. D4693-1G) in HBSS at 37 °C for approximately 90 min with shaking at 25 rpm. DNase I (Sigma, Cat. No. 4716728001) was added during the final 10 min to prevent cell aggregation. The cell suspension was filtered through a 40 μm nylon cell strainer (Falcon, Cat. No. 352340). Dissociated single cells were washed 2–3 times with HBSS containing 0.1% (w/v) BSA (Gibco, Cat. No. 15260073) and diluted to approximately 100 cells/μL for downstream single-cell RNA sequencing.

### 10x Chromium single-cell library preparation and sequencing

Single-cell encapsulation and cDNA libraries were prepared using the Chromium™ Single Cell 3′ Reagent Kits v3 and Chromium™ Single Cell A Chip Kit (10x Genomics) according to the manufacturer’s protocol. Cells were loaded to capture 5,000–10,000 cells per chip position. All subsequent steps, including reverse transcription, cDNA amplification, and library construction, followed the standard 10x Genomics protocol. The final libraries were sequenced on an Illumina NovaSeq 6000 platform using 150 nt paired-end sequencing with a target of 100 GB raw reads per sample.

### Single-cell RNA sequencing data processing

Raw reads were processed using the Cell Ranger pipeline (10x Genomics) with default settings. FASTQ files were aligned to the human reference genome (GRCh38). Gene-barcode matrices were generated by counting unique molecular identifiers (UMIs) and filtering out non-cell-associated barcodes. The resulting gene-cell matrices were imported into the Seurat R toolkit (v5.5.0) for quality control and downstream analysis. Doublets were detected using scDblFinder with default parameters, which classified 13.27% of cells as predicted doublets. As a sensitivity analysis, clustering and cell-type annotation were repeated after excluding predicted doublets; all 12 annotated cell populations remained identifiable at closely comparable proportions, indicating that the cell-type annotations and composition are robust to doublet contamination.

### 10x Visium spatial transcriptomics

Fresh tumor tissue was embedded in optimal cutting temperature (O.C.T.) compound and frozen at −80 °C. Sections of 10 μm thickness were cut using a cryostat and placed onto a Visium Spatial Gene Expression slide. Tissue sections were fixed with pre-chilled methanol, stained with haematoxylin and eosin (H&E), and imaged using a brightfield slide scanner. Permeabilization time was optimized using the Visium Spatial Tissue Optimization Slide & Reagent Kit according to the manufacturer’s instructions.

After permeabilization, reverse transcription was performed on the slide to generate spatially barcoded cDNA. Second-strand cDNA synthesis was followed by denaturation and amplification. Libraries were constructed using the 10x Genomics Visium Spatial Gene Expression Reagent Kit according to the standard protocol. The resulting libraries were sequenced on an Illumina NovaSeq 6000 system. Raw sequencing data were processed using Space Ranger v3.0.0 (10x Genomics) with the GRCh38 human reference genome to generate gene-spot count matrices.

### Mitochondrial DNA variant enrichment and sequencing

To enrich mitochondrial transcripts from spatial transcriptomics libraries, we performed the MAESTER protocol as previously described (Miller et al. 2022; Xue et al. 2026). Briefly, whole-transcriptome amplification products from the 10x Visium spatial gene expression workflow were used as template for two rounds of PCR amplification with primer pools tiling the entire mitochondrial transcriptome. Each primer contained a 5′ handle for Illumina adapter addition. The first PCR (6 cycles) selectively amplified 15 mitochondrial transcripts. Products were pooled and purified with 1× AMPure XP beads. The second PCR (6 cycles) added full Illumina adapters (P5/P7) and dual-sample indices. Final libraries were purified with 0.8× AMPure XP beads and sequenced on an Illumina NovaSeq platform system with 28 cycles for Read 1, 8 cycles for i7, 8 cycles for i5, and 256 cycles for Read 2. This MAESTER enrichment enabled high-depth mitochondrial DNA sequencing from the same spatial spots used for transcriptomic profiling.

### RNA velocity analysis using cell2fate

To reconstruct the temporal progression of tumor cell states and infer transcriptional trajectories, we employed cell2fate (Aivazidis et al., 2025). Specifically, we applied the Cell2fate_DynamicalModel, which automatically determined the number of modules based on the underlying data structure, to capture transcriptional dynamics. The model was trained using a GPU-accelerated PyTorch backend with default parameters, running multiple iterations until convergence. Finally, the latent time inferred from RNA velocity and the module activation states were projected onto the UMAP embedding to visualize the resulting developmental trajectories.

### Integration of RNA velocity modules with spatial transcriptomics

To link temporal transcriptional programs to tissue architecture, we mapped the cell2fate modules onto the 10x Visium spatial transcriptomics data. Using the steady-state expression signatures of each module (i.e., the expected spliced counts at *t* → ∞) as reference profiles, we applied cell2location (Kleshchevnikov et al. 2022) to deconvolve the abundance of each module across Visium spots. Briefly, cell2location uses a Bayesian model to estimate the absolute contribution of each reference signature per spot, accounting for spot-level total RNA counts and technical variation. This approach yielded spatial maps of module activity, revealing where specific transcriptional programs (e.g., proliferative, immunosuppressive, pre-metastatic) are active within the tumor tissue.

### Spatial clustering and pseudotime-space trajectory inference (stlearn)

The sequencing data and imaging TIFF files from 10x Visium slides were processed using SpaceRanger software aligned to the GRCh38 human reference genome. The resulting count matrices were loaded into the stlearn in Python (Pham et al. 2023). A spatial graph was constructed by connecting physically adjacent Visium spots. Louvain clustering was performed on the spatial graph, and spatially non-contiguous clusters were further split by sub-clustering to obtain five final spatially coherent clusters (Clusters 0 to 4).

To reconstruct transcriptional progression, we applied the pseudo-time-space (PSTS) algorithm. PSTS combines gene expression distance (cosine distance on PCA-reduced data) and spatial distance (Euclidean distance between spot centroids) into a weighted distance *d_PTS_* = *ω* ⋅ *d_PT_* + (1 − *ω*) ⋅ *d_S_* , with the weight *ω* optimized by graph spectral comparison. A minimum spanning arborescence was computed on the directed graph of sub-clusters to infer spatial trajectories from COL3A1⁺ progenitors toward ALPL⁺ and THY1⁺ lineages.

Transition genes along each branch were identified by Spearman correlation of gene expression with PSTS values (|ρ| > 0.3, adjusted p < 0.05), followed by GO and KEGG enrichment analysis.

### Quantification of pseudotemporal progression, pathway activity, and spatial gene expression patterns

#### Pseudotime distribution and differential expression analysis

For each spatial spot, a pseudotime value was assigned using the pseudo-time-space (PSTS) algorithm described above. Pseudotime distributions between clusters were compared using a two-sided Wilcoxon rank-sum test. Differentially expressed genes (DEGs) between clusters were identified using the scanpy.tl.rank_genes_groups function with the Wilcoxon test and Benjamini-Hochberg correction (adjusted p < 0.05).

#### Pathway activity scoring

To assess the activation of specific biological pathways in individual spatial spots, we performed single-sample pathway activity scoring using the gseapy Python package. For each spot, an enrichment score was calculated based on the average expression of curated gene signatures from MSigDB Hallmark and Reactome databases. Specifically, we evaluated the NF-κB pathway using genes from “HALLMARK_TNFA_SIGNALING_VIA_NFKB” and “NF-κB activation” (e.g., *NFKB1*, *HLA-B*, *CD74*, *TMSB4X*); fatty acid metabolism using “HALLMARK_FATTY_ACID_METABOLISM” (e.g., *ACSL1*, *CPT1A*, *FASN*); EMT using a signature of *VIM*, *CDH2*, *SNAI1*, *TWIST1*, *ZEB1*; and cell cycle (proliferation) using G2/M phase genes (*TOP2A*, *KIF20A*, *CCNB1*, *AURKB*). Pathway scores were compared between clusters (e.g., Cluster 1 vs. 2 or Cluster 1 vs. 4) using a two-sided Wilcoxon rank-sum test (adjusted p < 0.05). Additionally, Spearman correlation was used to assess the relationship between each pathway score and pseudotime along a given trajectory.

#### Trajectory-correlated gene expression and EMT scoring

For each gene, the Spearman correlation between its log-normalized expression and pseudotime was computed on spots pooled across both clusters of each transition (Cluster 1 and Cluster 2 for the 1 to 2 transition; Cluster 1 and Cluster 4 for the 1 to 4 transition), with cluster identity indicated by dot color in the scatter plots. Genes with |ρ| > 0.2 and adjusted p < 0.001 were considered significantly correlated.

The EMT score per spot was calculated as the average expression of a predefined EMT gene signature (VIM, CDH2, SNAI1, TWIST1, ZEB1). Scores were plotted against pseudotime, and local regression (LOESS) was used to visualize the trend. Spots from different clusters were color-coded.

#### Spatial mapping of gene expression

Normalized expression values of selected genes (VIM, CALD1, COL1A1, MALAT1) were visualized on the tissue section using the spatial coordinates provided by the Visium platform. Each spot was colored according to its expression level, and cluster identities were overlaid to assess spatial enrichment.

#### Functional enrichment analysis

Differentially expressed genes between clusters were subjected to Gene Ontology (GO) biological process and KEGG pathway enrichment analysis using the gseapy library (Enrichr) or clusterProfiler (R). Significance was defined by an adjusted p-value < 0.05 (Benjamini-Hochberg). The top enriched terms were visualized as bar plots.

#### Cell-cell interaction analysis using CellChat V2 with spatial constraints

To infer ligand-receptor mediated cell-cell interactions, we used CellChat V2.1.0 (Jin et al. 2025). A CellChat object was created from the Seurat object containing spatial coordinates, spot-level gene expression, and cell type annotations (dominant cell type per spot). The ‘trimean’ parameter was enabled to filter out weak (noise) interactions. Only interactions between spots within a Euclidean distance ≤250 μm were retained to model paracrine signaling, consistent with the typical molecular diffusion range. For each signaling pathway, we computed global communication probabilities and ranked ligand-receptor pairs by their relative contribution. Spatial interaction patterns were visualized using the CellChat spatial projection function.

#### SpaCET-based deconvolution, malignancy inference, and spatial CCI

As an orthogonal approach, we applied SpaCET V1.3.0 (Ru et al. 2023), a spatial-transcriptomics-native tool, to the same Visium dataset. A SpaCET object was created directly from the Space Ranger output using create.SpaCET.object.10X(). Quality control was performed with SpaCET.quality.control(), filtering spots with <100 detected genes. Cell type deconvolution was carried out using SpaCET.deconvolution() with the cancer type set to “SARC” (sarcoma), leveraging SpaCET’s internal reference database of 30 solid tumor types. Malignant cells were identified by copy number alteration (CNA) inference; further, malignant cells were stratified into “Malignant State A” (less aggressive) and “Malignant State B” (more aggressive) based on hierarchical clustering of CNA and gene expression signatures. Spatial tumor-stromal interfaces were identified using SpaCET.identify.interface().

#### Identification of informative mitochondrial DNA variants

Raw sequencing reads were aligned to the human reference genome (GRCh38) using Cell Ranger. For phylogeny construction with MitoDrift, we followed the variant calling, filtering, and quality control procedures as described in the original MAESTER publication (Miller et al. 2022). This pipeline yielded 5,263 informative mitochondrial variants across 4,888 spots for phylogenetic reconstruction, following the variant-selection recommendations of the MitoDrift paper (T. Gao et al. 2026)

#### Phylogenetic tree reconstruction and lineage distance

To infer the lineage history of tumor cells based on mitochondrial variants, we reconstructed a phylogenetic tree using MitoDrift (T. Gao et al. 2026), a Wright-Fisher drift model coupled with a hidden Markov tree (WF-HMT). This approach models the stochastic inheritance and drift of heteroplasmic mitochondrial variants across cell divisions. The hidden Markov tree captures the hierarchical relationships among cells, with transition probabilities governed by the Wright-Fisher process. The MitoDrift phylogenetic reconstruction used 5,263 informative mitochondrial variants across 4,888 spots, filtered according to the criteria described in the original MAESTER and MitoDrift publications. This variant set is essentially non-overlapping with the 7 spatially autocorrelated variants used for vireoSNP clonal deconvolution (Fig. 6); only one of the seven (709G>A) is also present in the MitoDrift variant set. Posterior clade support was estimated, and low-confidence branches were collapsed using a threshold of τ = 0.125. The resulting tree comprised clades defined by unique combinations of mitochondrial variants, where each spot (or cell) was annotated with its predominant cell type inferred from cell2location deconvolution.

Phylogenetic uncertainty was quantified with 16 independent MCMC chains of 10,000 iterations each (2,000-iteration burn-in; 128,016 post-burn-in samples), within the 10–50-chain range recommended by the MitoDrift authors. MCMC convergence was assessed using the average standard deviation of split frequencies (ASDSF), which reached a stable plateau of 0.320 (flat over the final 3,000 iterations). This plateau reflects the challenge of achieving full topological convergence across all bipartitions in a tree with 4,888 tips, where rare shallow splits contribute disproportionately to the metric. Per-chain concordance analysis (Fig. S10) provides a more granular assessment: 100% of clades with posterior support ≥ 0.80 (n = 451) are recovered in at least 12 of 16 independent chains. Model parameters (ngen = 1.016, ε = 0.00242, err = 0.0148) were estimated from the same input data by expectation–maximization; an independent replication run with parameter re-fitting enabled reproduced all three estimates exactly and yielded byte-identical MCMC output. The phylogeny is rooted by construction: MitoDrift introduces a dummy outgroup tip with all-zero variant allele frequency, representing the ancestral (mutation-free) mtDNA state; the tree is rooted on this outgroup and the outgroup is then dropped, fixing the root state to VAF ≈ 0 (ancestral).

Ancestral cell states were reconstructed by maximum parsimony on the rooted tree, with DELTRAN resolution favouring parallel gains; robustness of the inferred root state was assessed by re-rooting under midpoint and burden-minimal strategies. Derived-variant burden per spot (number of variants with VAF ≥ 0.1) was residualised on sequencing depth (log1p burden ∼ log depth) for model-free ordering analyses.

For each pair of spots, the phylogenetic distance was calculated as the cophenetic distance on the WF-HMT tree. Transcriptomic distance was computed as the Euclidean distance between spots based on the first 30 principal components of gene expression.

Mutation–clade association screen. To determine whether the inferred clades are supported by distinct mitochondrial variants, each informative variant was tested for association with every high-purity clade. For each of the 255 maximal high-purity single-state clades (purity ≥ 0.8, ≥2 tips) and the three clades highlighted in Fig. 7, variant presence (VAF ≥ 0.1) within the clade was compared with all other tips using a one-sided Fisher exact test, with Benjamini–Hochberg correction across all 53,159 variant×clade tests. Complete results are provided in Supplementary Table 1.

#### VAF-based clonal deconvolution with spatial quality control

**(a) Input preparation.** Mitochondrial variant calls for the Visium-MAESTER dataset were generated with cellsnp-lite and filtered by MQuad (Kwok et al. 2022), yielding 8 candidate variants across 1,849 spots. Clone assignments were inferred with vireoSNP (Huang et al. 2019) using the BinomMixtureVB algorithm. Following the Moran’s I filter described in (b), 12136T>C was excluded and vireoSNP was re-run on the remaining 7 variants. The number of clones was selected by a BIC sweep over n_clones ∈ {4..10} (n_init = 10, random_state = 42), yielding 6 clones. Analyses were performed in Python 3.9 with numpy ≥ 1.24 and vireoSNP 0.5.9.
**(b) Per-variant Moran’s I and multiple-testing correction.** Spatial autocorrelation of each variant’s allele frequency was quantified with Moran’s I,

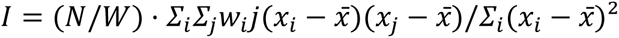

where the indices i, j run over the N spots with per-variant depth DP ≥ 5, x_i is the allele frequency at spot i, and ̄x is its mean across the N valid spots. The spatial weight w_ij is a binary indicator that takes the value 1 when spots i and j are hex-adjacent on the Visium array — that is, when their Euclidean distance in (array_row, array_col) index units is less than r = 2.1 — and 0 otherwise; by construction w_ii = 0 and w_ij = w_ji. The normalisation constant W = Σ_i Σ_j w_ij is therefore equal to twice the number of unordered hex-adjacent pairs (7,057 such pairs on the B82 section, so W = 14,114). Pairs with either endpoint marked invalid for a given variant were excluded from both the numerator and the denominator for that variant. The closed-form Z statistic, Z = (I_obs − μ_null) / σ_null, was computed from the mean and standard deviation of I under a label-shuffle null in which the VAF values were permuted among the N valid spots; we then derived a normal-approximation p value from Z and, for robustness, also reported the one-sided empirical p value p_perm = (1 + #{I_null ≥ I_obs}) / (1 + B) computed across B = 9,999 permutations (seed = 42). Note that the permutation null is non-parametric, while the closed-form Z and its normal p value assume only that I under the null is approximately Gaussian — a different assumption from the variance-component models used by SpatialDE (Svensson et al. 2018), SPARK (Sun et al. 2020) and SUMMIT (Bracht et al. 2025), and one that is appropriate for use as a variant-quality filter on a small candidate set of mitochondrial variants. Multiple testing across the 8 candidate variants was controlled by a Bonferroni-corrected threshold α = 0.05 / 8 = 0.00625.

**(c) Depth-autocorrelation control.** Moran’s I of per-spot total depth (summed across the 8 candidate variants) was computed on the same hex-neighbour graph and under the same 9,999-permutation null. When the depth Z exceeded a conservative threshold of 3.0 (corresponding to a two-sided normal-approximation p ≈ 0.003, indicating that sequencing depth is itself non-randomly distributed across the section), each variant’s allele frequency was residualised against per-spot total depth by ordinary least squares and Moran’s I was recomputed on the residuals.
**(d) Clone spatial coherence.** The fraction of hex-adjacent spot pairs assigned to the same clone (across the 7,057 hex pairs) was compared to a label-shuffle null in which the clone labels were permuted across the 1,849 spots (B = 9,999 permutations, seed = 42). The Z statistic Z = (f_obs − μ_null) / σ_null was converted to a one-sided upper-tail p value via the normal-approximation survival function (scipy.stats.norm.sf); the empirical permutation p value was reported in parallel as (1 + #{f_null ≥ f_obs}) / (1 + B). A negative control was generated by permuting the (row, col) coordinates of the spots and recomputing the same statistic on the unchanged clone labels.
**(e) Indirect strand-bias check.** Per-strand cellSNP output was not preserved for the B82 dataset, precluding a direct strand-balance test. As an indirect substitute, we computed the substitution-class distribution across all 16,192 MQuad candidate variants in the chrM region: >A substitutions accounted for 49.2% of candidates versus the 25% expected under a uniform substitution model. Six of the seven retained variants lay within a 12-nt window spanning positions 1861–1872 (inclusive) and were all substitutions to A — a pattern consistent with, though not definitive of, a strand-specific chemistry artefact in that region (Fig. S7A). The only retained variant outside this window, 709G>A, carries the second-strongest Moran’s I after 1872T>A and passes the depth-residualization control independently. To test directly whether the spatial clone signal depends on this window, we re-ran the full vireoSNP clone-calling (BIC sweep over n_clones ∈ {4..10}, n_init = 10, seed = 42) and the hex-neighbor same-clone spatial-coherence permutation test (9,999 label shuffles, seed = 42) under three leave-out variant sets: (i) dropping only 1872T>A (six variants; Z = 11.0); (ii) dropping the entire 1861–1872 window so that only 709G>A remained (carrier vs. non-carrier split, 536 carrier spots; Z = 10.5); and (iii) the six in-window variants alone (Z = 9.4). All conditions remained significant at p < 10⁻⁴ (Supplementary Fig. S7B), demonstrating that the spatial clone result is not driven by the potentially strand-biased window.

#### Mantel test for lineage-state coupling

To test whether mitochondrial lineage history correlates with transcriptional divergence, we performed a Mantel test (999 permutations) (Mantel, 1967). between the phylogenetic distance matrix and the transcriptomic Euclidean distance matrix using the vegan package in R. A significant positive correlation indicates that cells sharing recent mitochondrial ancestry also exhibit similar transcriptional states. To control for spatial autocorrelation, we additionally performed a partial Mantel test conditioning on spatial distance, and we compared expression distances between each spot’s closest phylogenetic relative (sister lineage) and spatially matched null pairs using the Wilcoxon signed-rank test.

#### Phylogenetic signal (Blomberg’s K)

To assess whether cell type states are phylogenetically heritable, we calculated Blomberg’s K for each binary cell-type indicator (COL3A1⁺, ALPL⁺, THY1⁺) on the WF-HMT tree. Values close to 0 indicate no phylogenetic clustering (i.e., the trait is not conserved across the tree), whereas values near 1 indicate strong signal. Significance was assessed by comparing observed K to null distributions from 999 random permutations of tip states.

## Supporting information

Systematic Fisher-exact screen of mtDNA variants across high-purity phylogenetic clades.

## Author Contributions

Y. X. and Z. S. contributed equally to this work. Y. X., Z. S., K. S. C. C. and J. W.K. H. conceived the study. Y. X. and Z. S. conducted experiments and generated the data. Y. X. and Z. S. conducted most of the informatics analysis with the guidance of J. W.K. Y. X., Z. S., J. S., A. S. H.Y. C., L.W., M. C., J. P. Y. C and K. S. C. C. participated in the informatics analysis and discussion. Y. X., Z. S., K. S. C. C and J. W.K. H. wrote the manuscript. All authors edited the manuscript.

## Acknowledgements

The authors thank Dr. Peter van Galen at Harvard Medical School for generously sharing bioanalyzer-positive library controls and for critical discussions on this project. The authors thank CPOS at HKUMed for technical assistance with 10x Chromium, 10x Visium, and NovaSeq. This study is supported in part by GRF 17123223 (J.W.K.H) from the Hong Kong Research Grants Council, the Guangdong Natural Science Foundation (2023A1515011265), and the Seed Fund of The University of Hong Kong (2401103785).

---

**Fig. S1.**
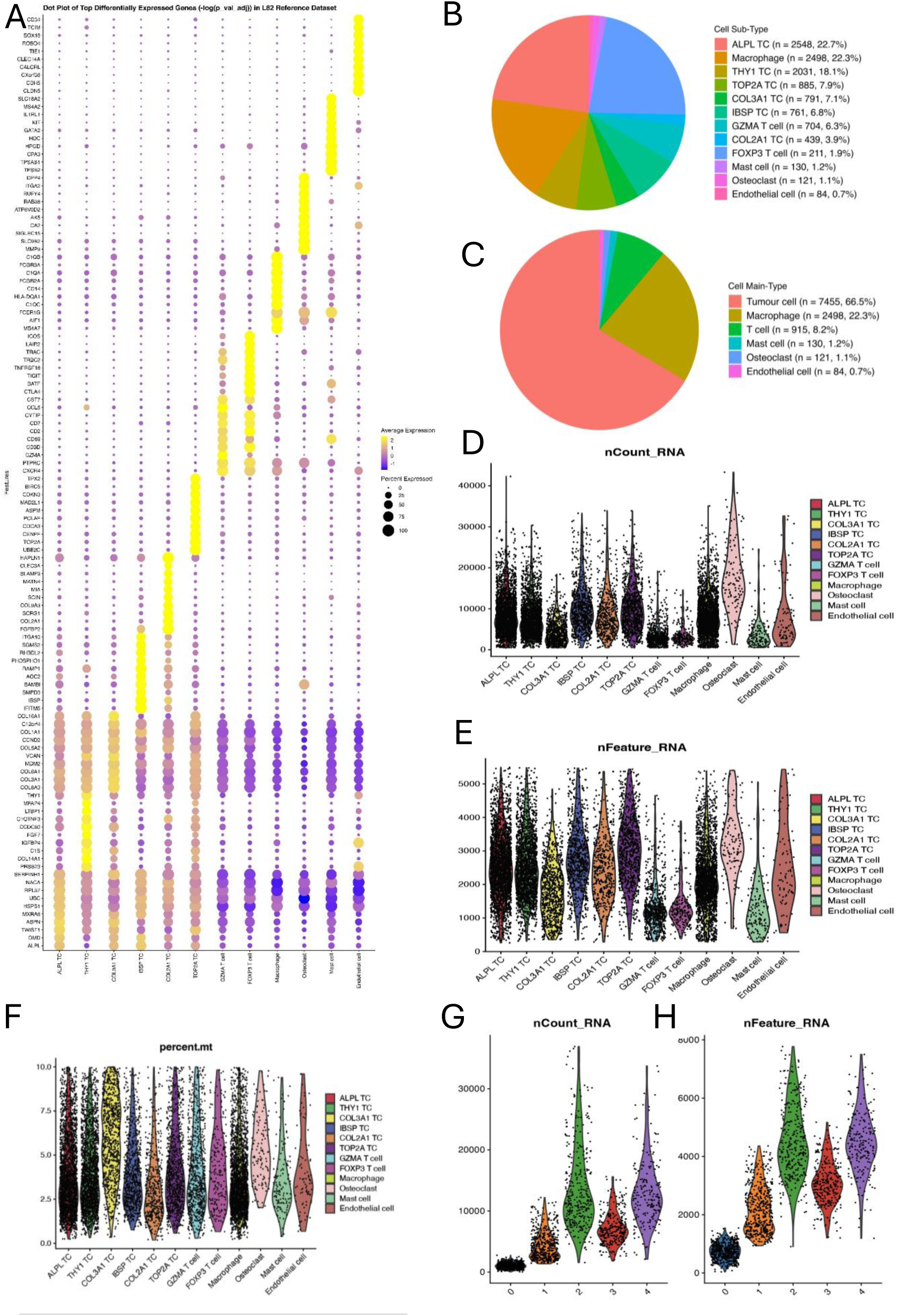
Analysis and quality control of single cell RNA-seq (A-F) and Visium spatial transcriptomics (G-H) data of osteosarcoma. (A) Dot plot showing the top marker gene of each cell subtype (B) Pie charts showing proportions of cell subtypes. (C) Pie charts showing proportions of major cell types. (D-F)Violin plots showing the distributions of (D) total unique molecular identifier (UMI) counts per cell (nCount_RNA), (E) the number of detected genes per cell (nFeature_RNA), and (F) the percentage of counts assigned to mitochondrial genes per cell (percent.mt) across the 12 annotated cell populations. (G-H) QC for Visium. Violin plots show the distributions of (G) total unique molecular identifier (UMI) counts per spot (nCount_RNA), (H) the number of detected genes per spot (nFeature_RNA).

**Fig. S2.**
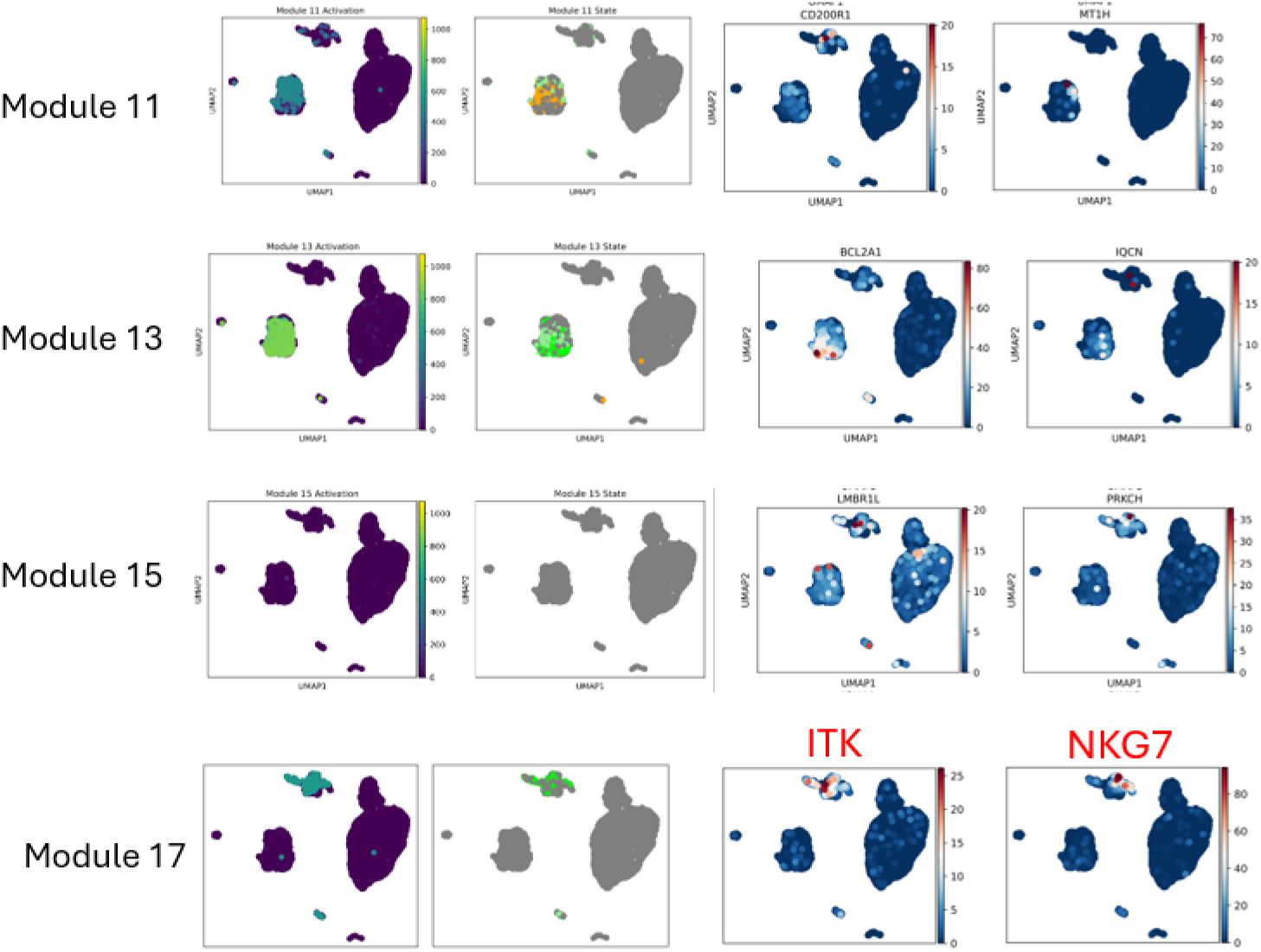
Gene module activation and marker gene expression across the tumor cell trajectory. UMAP visualizations showing the activation state of gene modules 11, 13, 15, and 17, along with representative marker genes for each module.

**Fig. S3.**
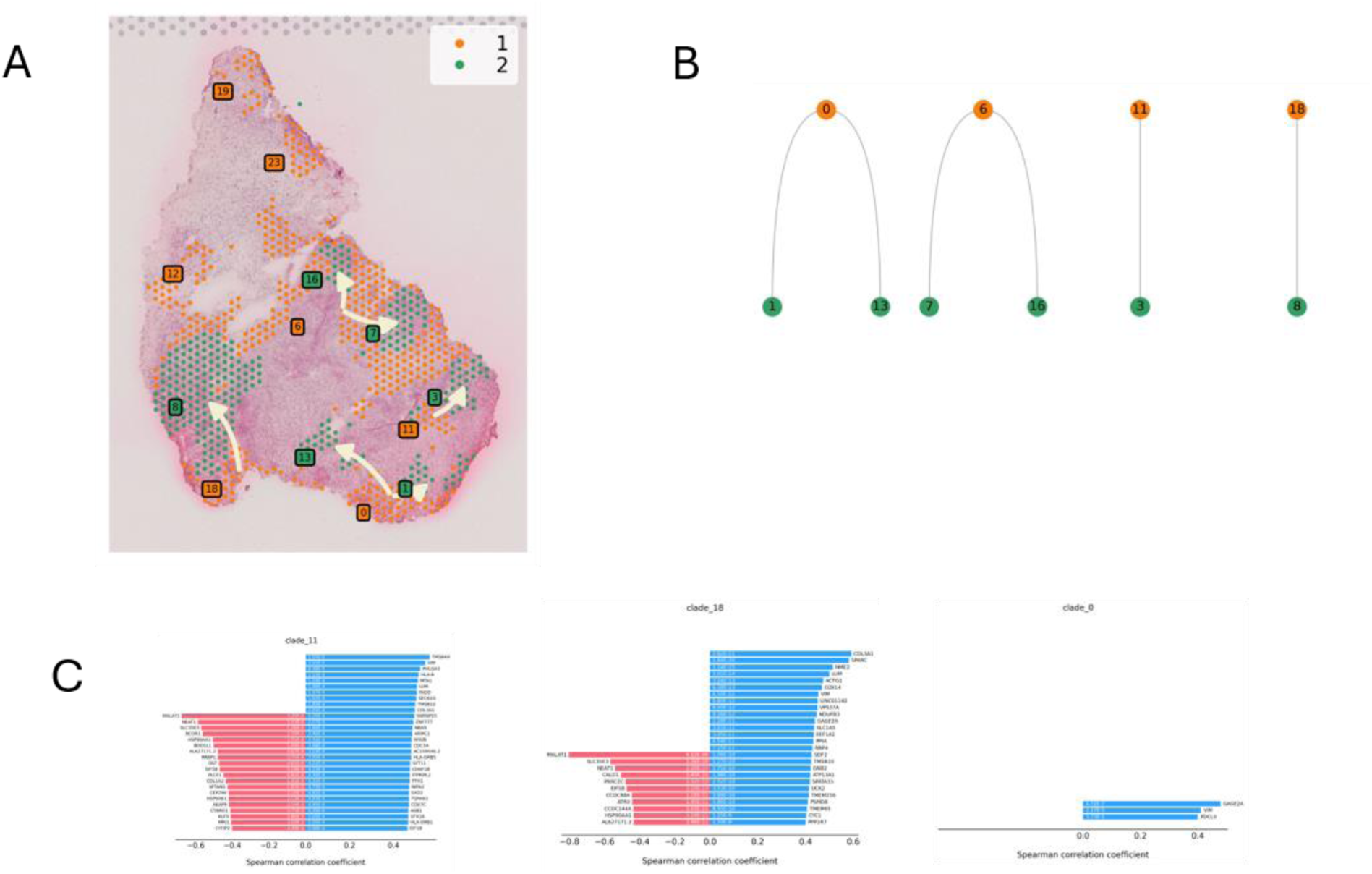
Molecular and spatial features of the pseudotime trajectory from Cluster 1 to Cluster 2. (A) Spatial distribution of Cluster 1 (orange) and Cluster 2 (green) cells in the osteosarcoma tissue section, with arrows indicating the inferred direction of cell state progression. (B) Schematic representation of the pseudotime trajectory connecting Cluster 1 and Cluster 2. (C) Spearman correlation coefficients showing gene expression dynamics across three clades along the trajectory (clade_18, clade_11, clade_0). Red bars indicate negative correlations and blue bars indicate positive correlations with pseudotime progression.

**Fig. S4.**
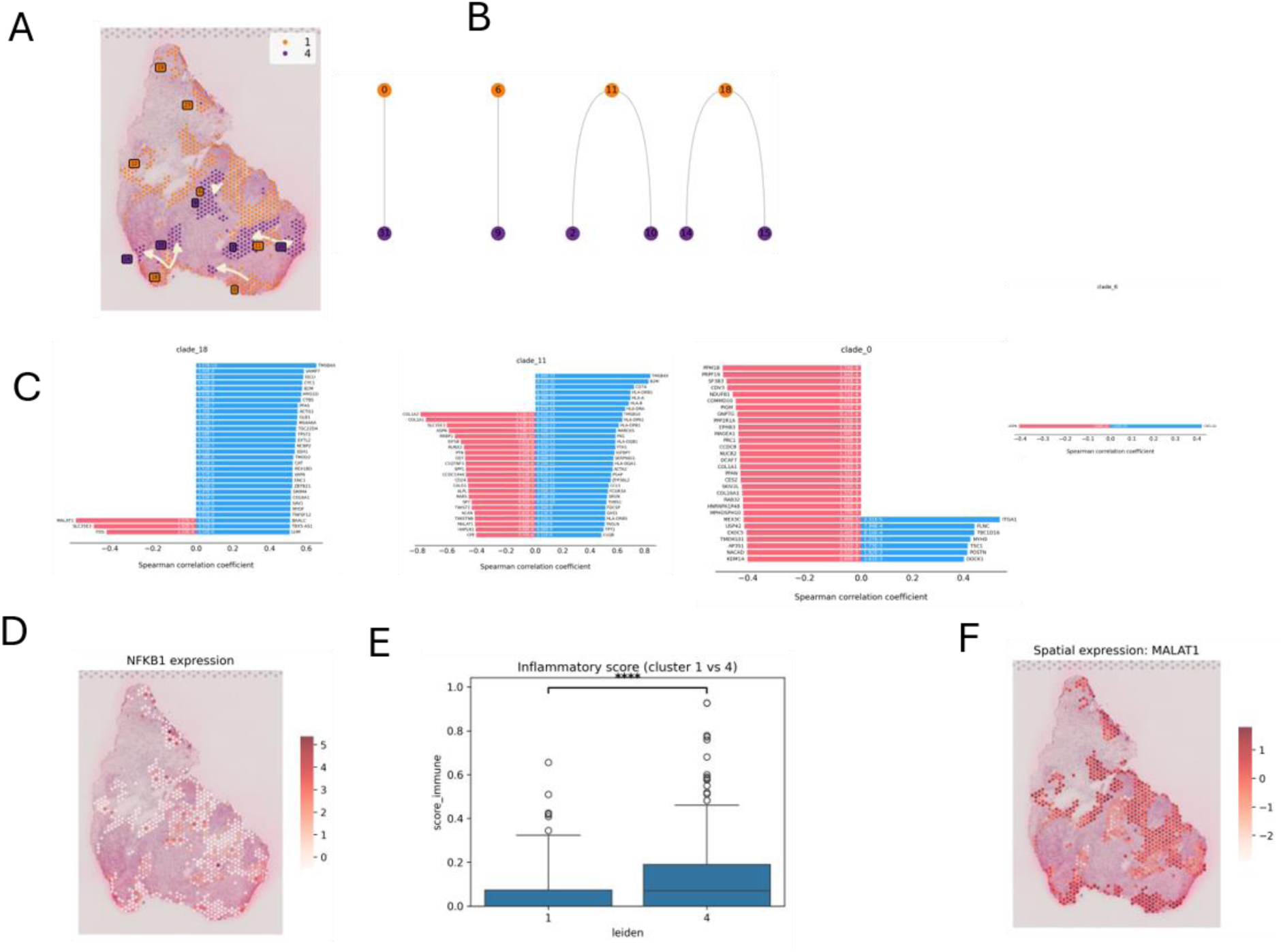
Molecular and spatial features of the pseudotime trajectory from Cluster 1 to Cluster 4. (A) Spatial distribution of Cluster 1 (orange) and Cluster 4 (purple) cells in the osteosarcoma tissue section, with arrows indicating the inferred direction of cell state progression. (B) Schematic representation of the pseudotime trajectory connecting Cluster 1 and Cluster 4. (C) Spearman correlation coefficients showing gene expression dynamics across three clades along the trajectory (clade_18, clade_11, clade_0). Red bars indicate negative correlations and blue bars indicate positive correlations with pseudotime progression. (D) Spatial expression patterns of *NFKB1* in the tissue section, revealing their differential distribution along the cluster 1-to-4 trajectory. (E) The inflammatory score is significantly higher in the THY1⁺ cluster (Cluster 4) than in the COL3A1⁺ cluster (Cluster 1). (F) Spatial expression patterns of *MALAT1* in the tissue section, revealing their differential distribution along the cluster 1-to-4 trajectory.

**Fig. S5.**
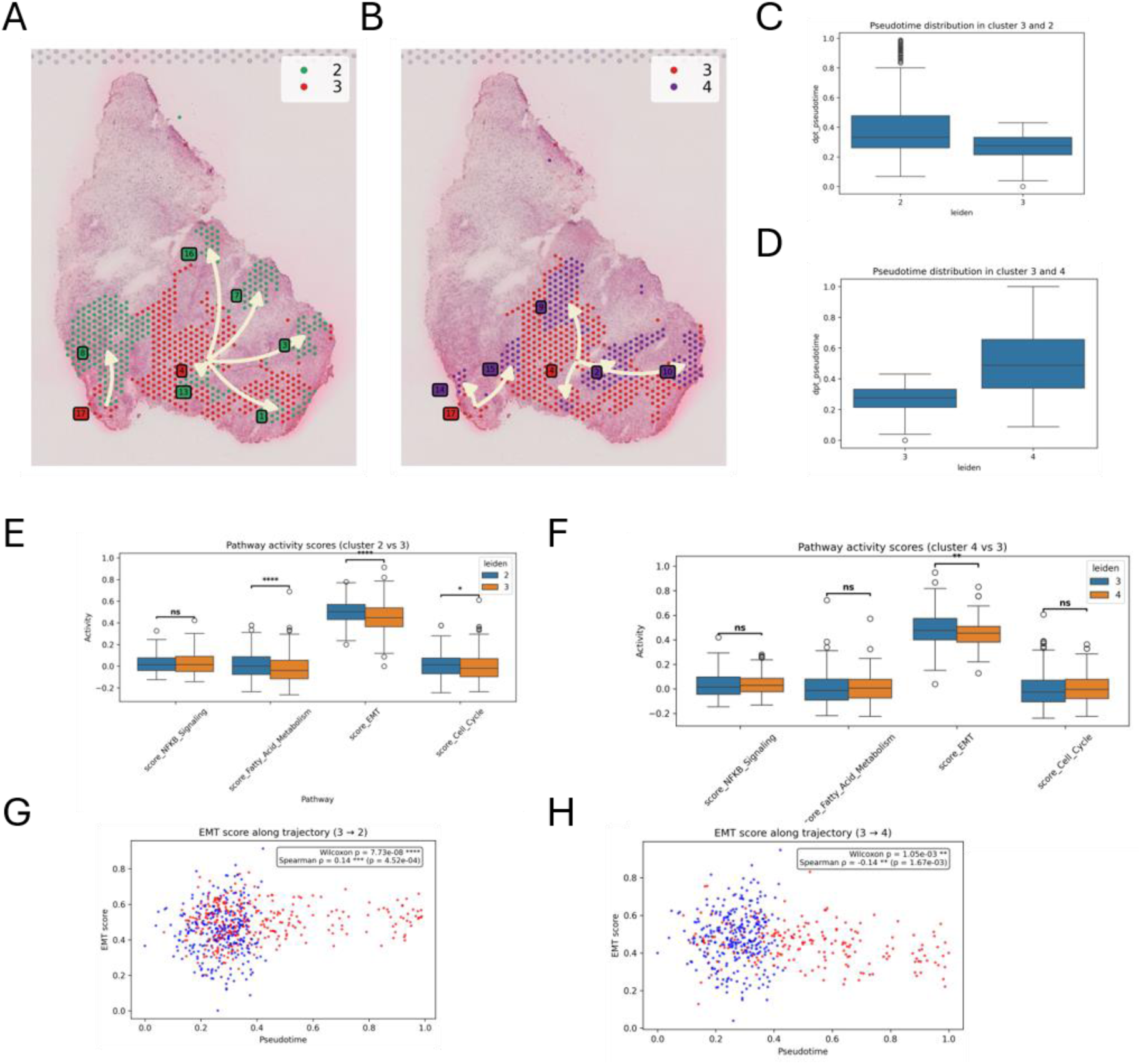
Spatial and functional characterization of cell state transitions along pseudotime trajectories in osteosarcoma. (A, B) Spatial distribution and pseudotime trajectories: 3 to 2 (A) and 3 to 4 (B). Arrows indicate the inferred direction of cell state progression. (C, D) Pseudotime distribution for the two trajectories. (E, F) Pathway activity scores comparing NF-κB, fatty acid metabolism, EMT, and cell cycle pathways. (G, H) EMT score dynamics along the trajectories with Spearman correlation analysis.

**Fig. S6.**
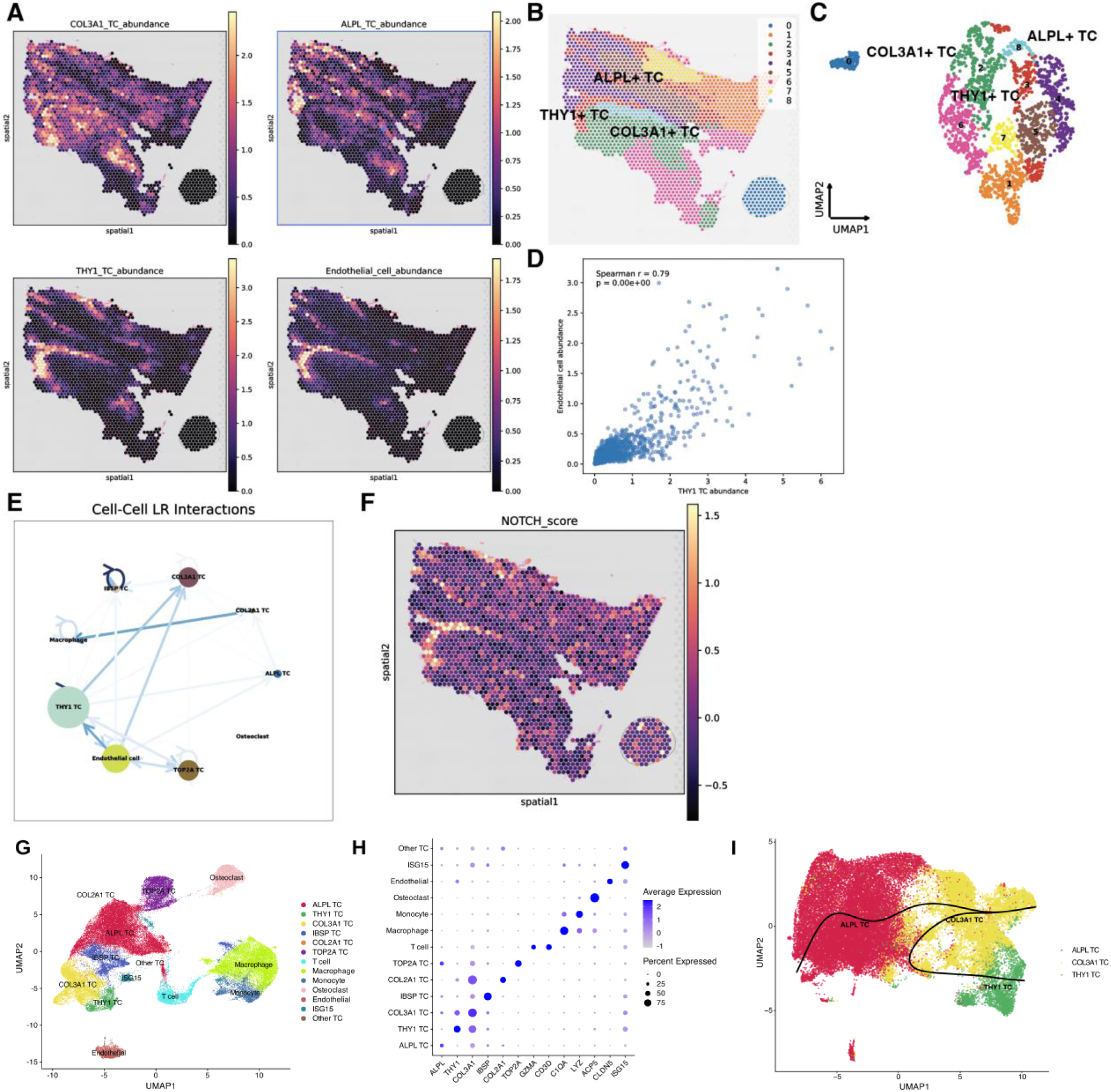
External validation of tumor-cell states and the THY1_Endo niche in independent osteosarcoma cohorts. (A) Spatial maps of cell2location-inferred abundances of COL3A1⁺, ALPL⁺, and THY1⁺ tumor cell states and endothelial cells. (B) Spatial distribution of spatial clusters. (C) UMAP of Visium spots, colored by cluster as in panel B. (D) Association between the inferred abundances of THY1⁺ tumor cells and endothelial cells. (E) Inferred ligand–receptor communication network. Nodes represent annotated cell populations, and directed edges represent inferred communication. (F) Spatial distribution of NOTCH-associated interaction scores. (G) UMAP of an independent osteosarcoma scRNA-seq cohort (GSE152048), coloured by annotated cell population; the COL3A1⁺, ALPL⁺ and THY1⁺ tumour-cell states are all recovered. (H) Dot plot of state-marker expression (colour: average expression; size: percent expressed) across the GSE152048 populations, validating the three tumour-cell states. (I) Slingshot trajectory inference on the COL3A1⁺, ALPL⁺ and THY1⁺ tumour cells of GSE152048, recovering COL3A1⁺ to ALPL⁺ and COL3A1⁺ to THY1⁺ lineages, consistent with the bifurcation observed in our cohort.

**Fig. S7.**
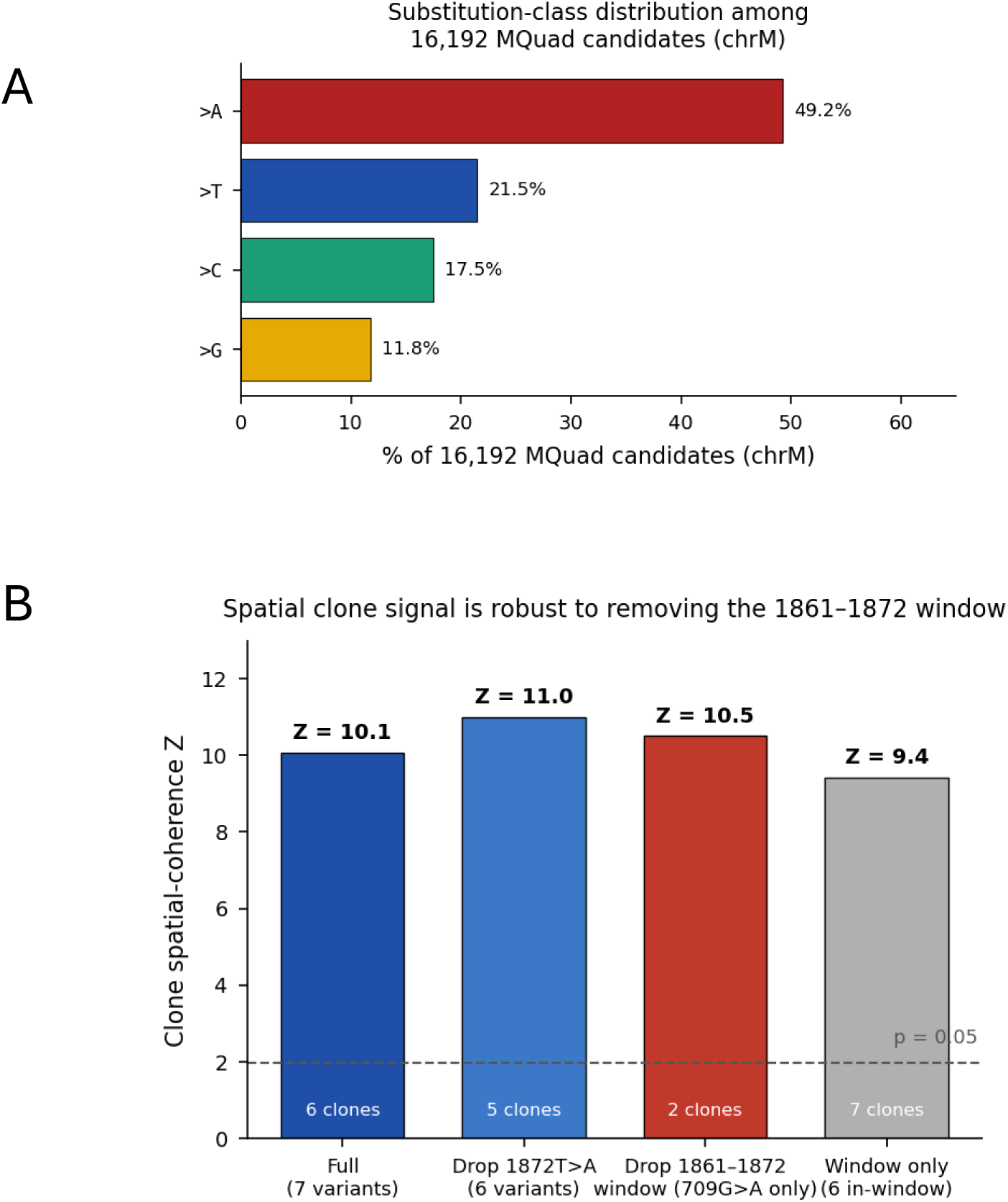
Strand-bias assessment and leave-out analysis of mtDNA spatial clone signal. (A) Substitution-class distribution among the 16,192 MQuad candidate variants in chrM. The >A class accounts for 49.2% of candidates versus the 25% expected under a uniform substitution model. Six of the seven retained variants lie within a 12-nt window spanning positions 1861–1872 (inclusive) and are all substitutions to A, consistent with — though not by itself proof of — a strand-specific chemistry artefact in that region; per-strand cellSNP output was not preserved, so a direct strand-balance test was not feasible. (B) Leave-out analysis of clone spatial coherence. Spatial-coherence Z after removing 1872T>A (Z = 11.0), removing the entire 1861–1872 window leaving only 709G>A (Z = 10.5), and using only the 1861–1872 window (Z = 9.4), compared with the full seven-variant set (Z = 10.1). Clone numbers are indicated within each bar. Dashed line: nominal significance threshold (Z = 1.96; p = 0.05); all four configurations also exceed the Bonferroni-corrected threshold (Z = 2.7; p = 0.00625). Coherence remains significant in all configurations, indicating that the spatial clone signal is not driven by the dominant variant, the 1861–1872 window, or any single site.

**Fig. S8.**
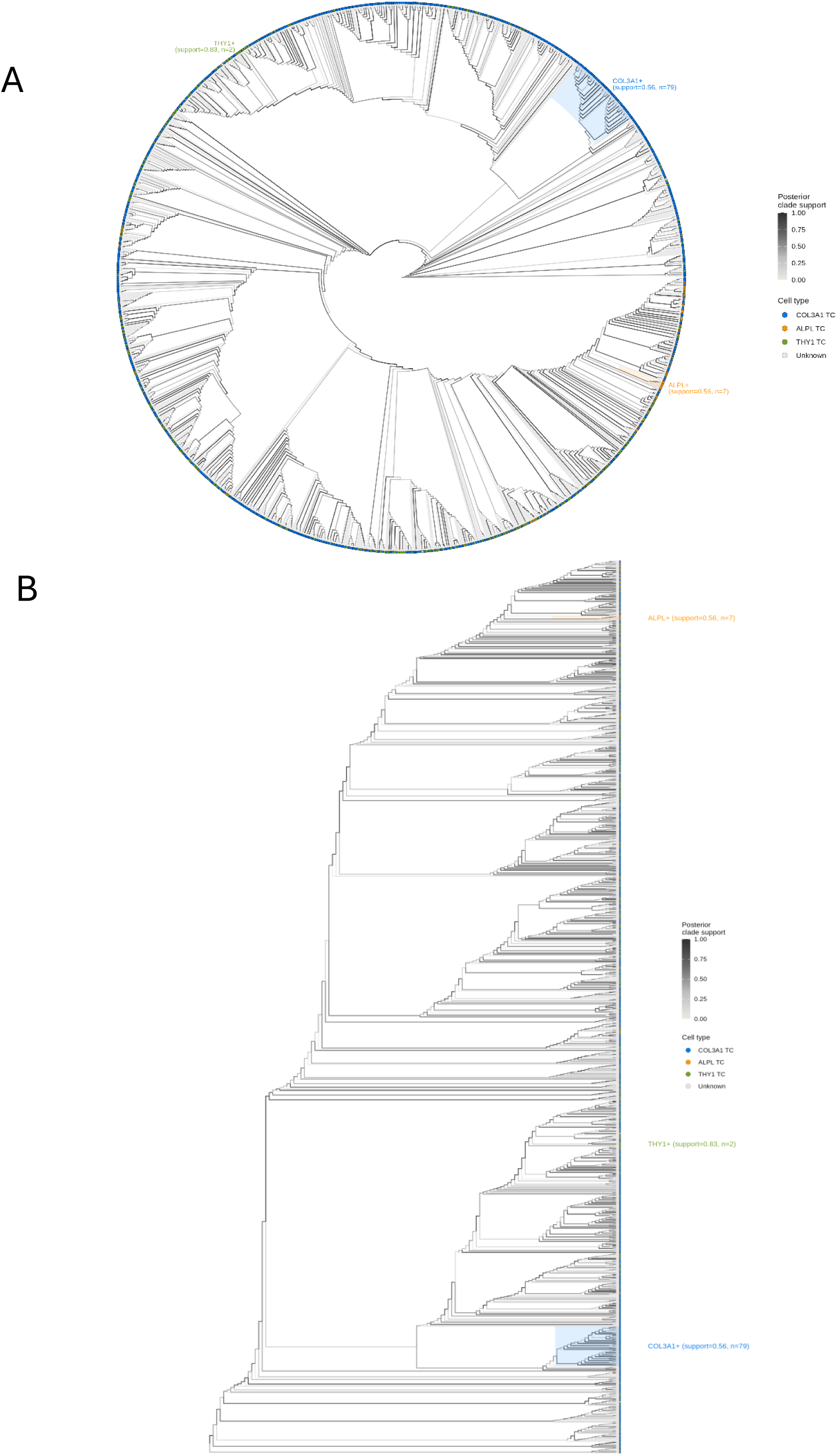
Posterior clade support mapped on the MitoDrift phylogeny. (A) Circular and (B) rectangular views of the confidence-refined mitochondrial phylogeny (inferred from 5,263 informative variants across 4,888 spots; low-confidence branches collapsed at τ = 0.125). Branches are coloured by posterior clade support computed from 16 independent MCMC chains (10,000 iterations each, 2,000-iteration burn-in); tip points are coloured by cell2location cell type (COL3A1⁺ = blue, ALPL⁺ = orange, THY1⁺ = green, unassigned = grey). The highest-purity clade for each tumour state is annotated with its posterior support and tip count.

**Fig. S9.**
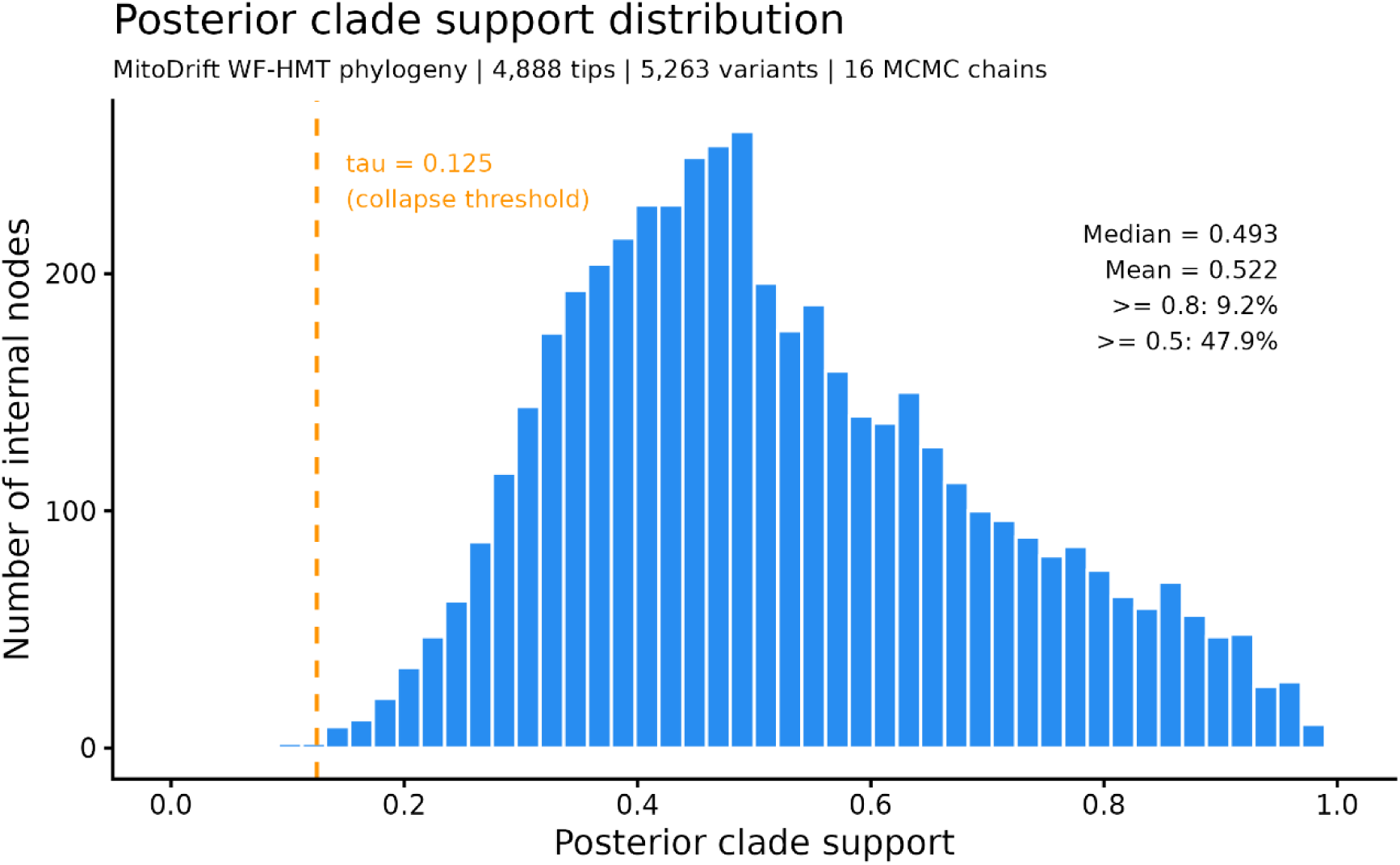
Distribution of posterior clade support. Histogram of posterior clade support across the 4,887 internal nodes of the phylogeny (16 MCMC chains). Median support = 0.493 (mean = 0.522); 47.9% of nodes have support ≥ 0.5 and 9.2% have support ≥ 0.8. The dashed line marks the τ = 0.125 collapse threshold used for display.

**Fig. S10.**
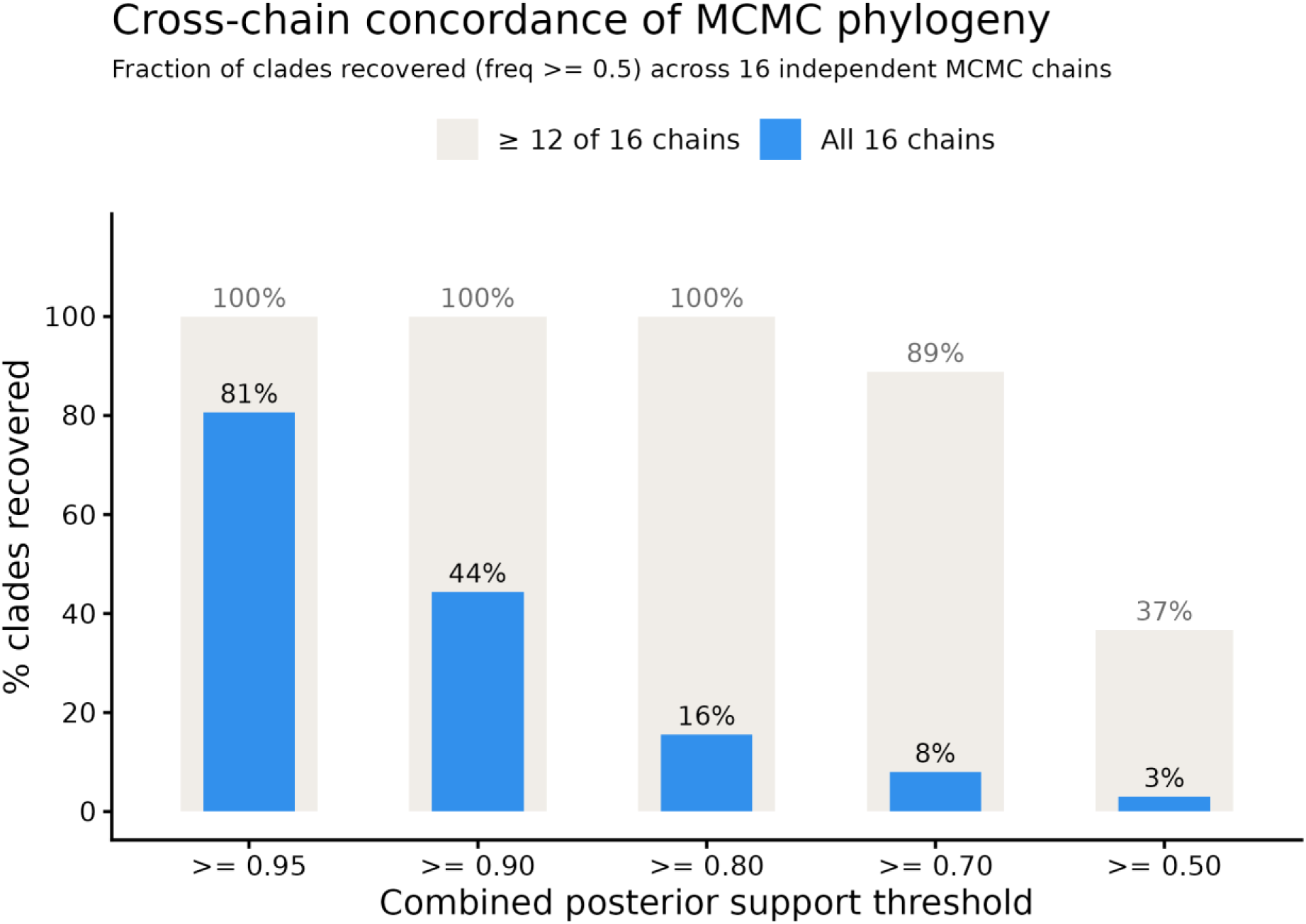
Cross-chain concordance of clade recovery. The 16 independent MCMC chains were treated as resampling replicates. For every internal node, the clade frequency was computed separately within each chain (8,001 post-burn-in samples per chain), and a clade was considered recovered in a chain at within-chain frequency ≥ 0.5. Bars show the percentage of clades, binned by combined posterior support, recovered in ≥ 12 of 16 chains (grey) or in all 16 chains (blue). All clades with combined support ≥ 0.80 (n = 451) are recovered in ≥ 12 of 16 chains.

**Fig. S11.**
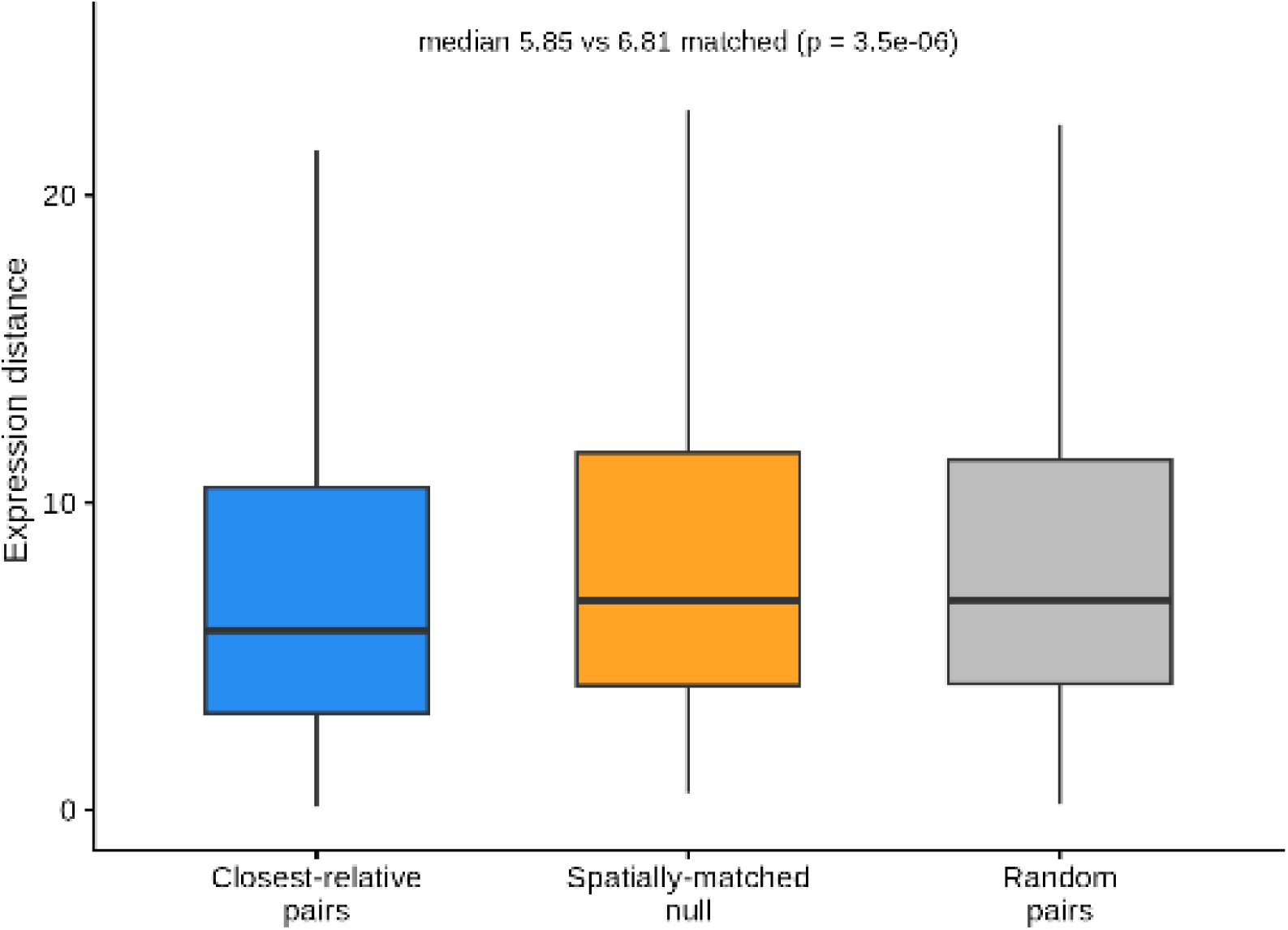
Sister-lineage test of lineage–state coupling. Expression distance between each spot and its closest relative on the tree (sister lineage; median 5.85) versus spatially matched null pairs (median 6.81; Wilcoxon signed-rank p = 3.5 × 10⁻⁶) and random pairs (median 6.83; p = 2.3 × 10⁻¹⁰).

**Fig. S12.**
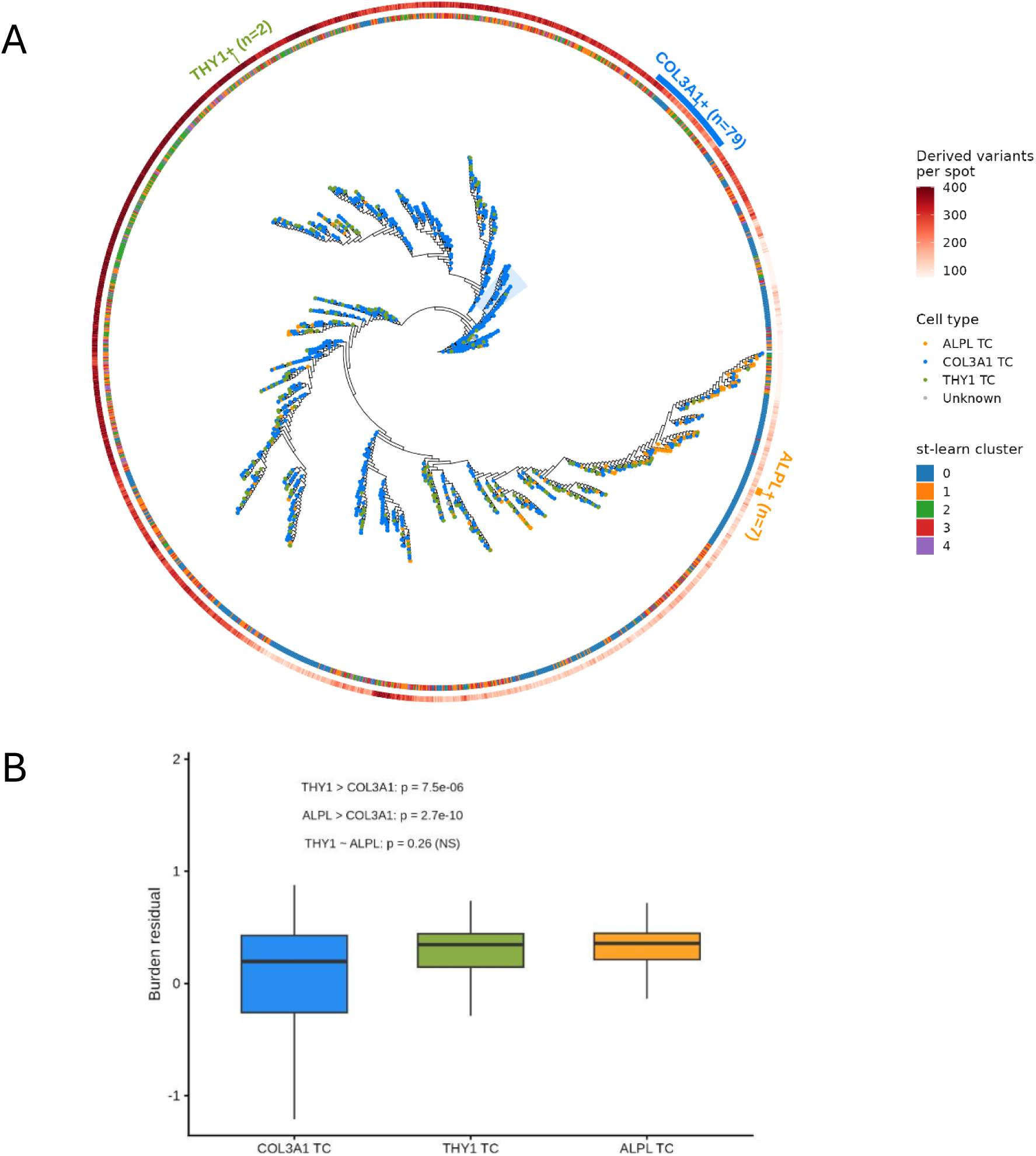
Per-spot derived-variant burden on the complete phylogeny. Circular cladogram of the cell2location-annotated phylogeny as in Fig. 7A, with a continuous outer ring encoding each spot’s derived-variant burden (number of variants with VAF ≥ 0.1; median 282, range 1–469).

**Fig. S13.**
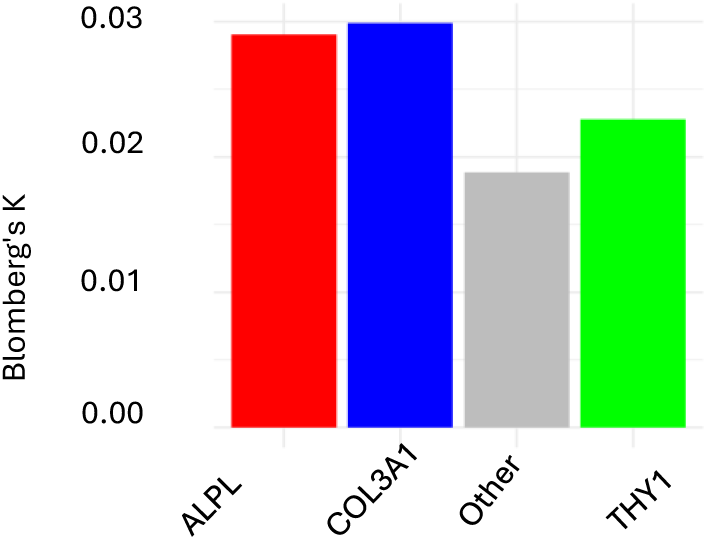
Blomberg’s K values for each cell type. Bar plot showing the phylogenetic signal (Blomberg’s K) for binary state-versus-rest indicators of the three tumour states on the cell-type-annotated subtree. K < 0.1 for all three states, indicating that none of the states is confined to a single monophyletic clade, consistent with repeated independent acquisition.

**Fig. S14.**
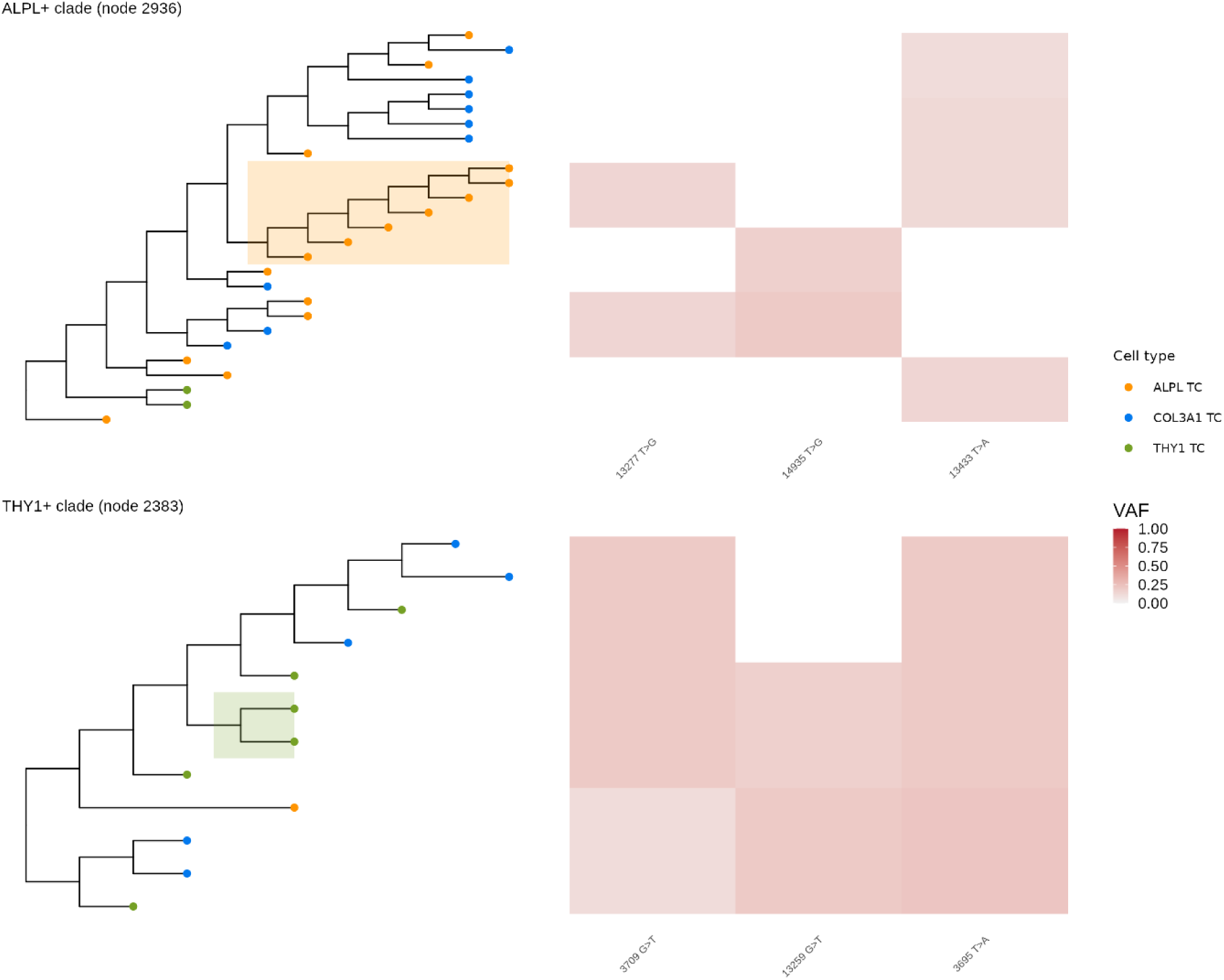
High-purity ALPL⁺ and THY1⁺ clades with per-spot VAF of clade-characteristic mutations. Focused subtrees as in Fig. 7C, with side panels showing the variant allele frequency (VAF) of the top clade-characteristic mtDNA mutations across the displayed spots.

**Fig. S15.**
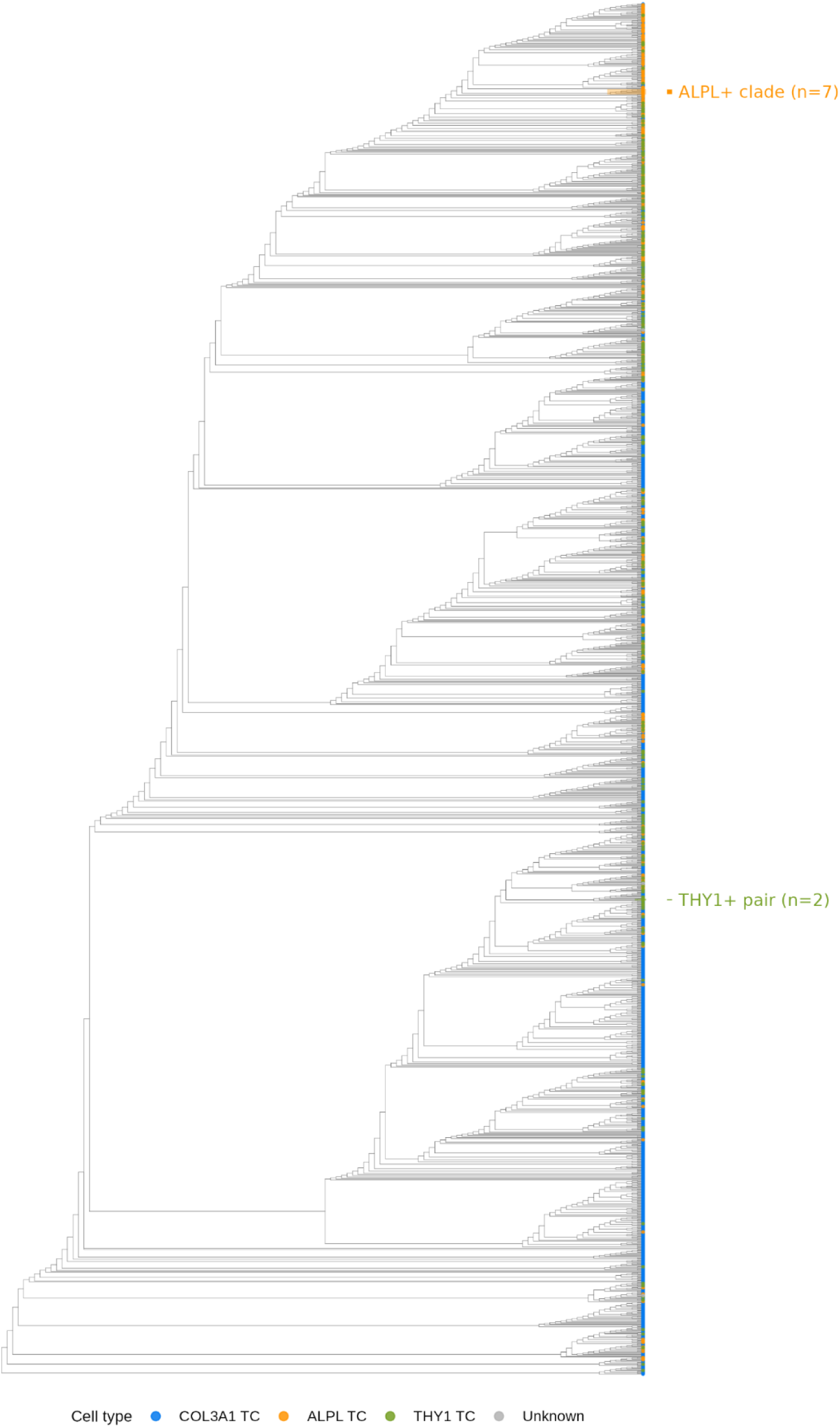
Rectangular view of the complete annotated phylogeny. Rectangular cladogram of the cell2location-annotated MitoDrift phylogeny as in Fig. 7A; tip points are coloured by cell type, and the high-purity ALPL⁺ (node 2936) and THY1⁺ (node 2383) clades are indicated.

